# The cryptic gonadotropin-releasing hormone neuronal system of human basal ganglia

**DOI:** 10.1101/2021.03.04.433872

**Authors:** Katalin Skrapits, Miklós Sárvári, Imre Farkas, Balázs Göcz, Szabolcs Takács, Éva Rumpler, Viktória Váczi, Csaba Vastagh, Gergely Rácz, András Matolcsy, Norbert Solymosi, Szilárd Póliska, Blanka Tóth, Ferenc Erdélyi, Gábor Szabó, Michael D. Culler, Cécile Allet, Ludovica Cotellessa, Vincent Prévot, Paolo Giacobini, Erik Hrabovszky

**Affiliations:** Laboratory of Reproductive Neurobiology, Institute of Experimental Medicine, 1083 Budapest, Hungary; Laboratory of Endocrine Neurobiology, Institute of Experimental Medicine, 1083 Budapest, Hungary; 1st Department of Pathology and Experimental Cancer Research, Semmelweis University, 1083 Budapest, Hungary; Centre for Bioinformatics, University of Veterinary Medicine, 1078 Budapest, Hungary; Department of Biochemistry and Molecular Biology, Faculty of Medicine, University of Debrecen, 4032 Debrecen, Hungary; Department of Inorganic and Analytical Chemistry, Budapest University of Technology and Economics, 1111 Budapest, Hungary; Department of Gene Technology and Developmental Biology, Institute of Experimental Medicine, 1083 Budapest, Hungary; Amolyt Pharma, Newton, MA, 02466 USA and 69-130 Ecully, France; Univ. Lille, Inserm, CHU Lille, U1172 - LilNCog - Lille Neuroscience & Cognition, F-59000 Lille, France

## Abstract

Human reproduction is controlled by ∼2,000 hypothalamic gonadotropin-releasing hormone (GnRH) neurons. Here we report the discovery and characterization of additional 150-200,000 GnRH-synthesizing cells in the human basal ganglia and basal forebrain. Extrahypothalamic GnRH neurons were cholinergic. Though undetectable in adult rodents, the GnRH-GFP transgene was expressed transiently by caudate-putamen cholinergic interneurons in newborn transgenic mice. In slice electrophysiological studies, GnRH inhibited these interneurons via GnRHR1 autoreceptors. Whole-transcriptome analysis of cholinergic interneurons and medium spiny projection neurons laser-microdissected from the human putamen confirmed selective expression of *GNRH1* and *GNRHR1* autoreceptors in cholinergic cells and uncovered the detailed transcriptome profile and molecular connectome of these two cell types. Higher-order non-reproductive functions regulated by GnRH under physiological conditions in the human basal ganglia and basal forebrain require clarification. GnRH/GnRHR1 signaling as a potential therapeutic target in the treatment of neurodegenerative disorders affecting cholinergic neurocircuitries, including Parkinson’s and Alzheimer’s diseases, needs to be explored.

## Introduction

Mammalian reproduction is controlled by a few hundred/thousand preoptic/hypothalamic neurons which release the decapeptide neurohormone gonadotropin-releasing hormone (GnRH) into the hypophysial portal circulation. GnRH promotes fertility via increasing the synthesis and secretion of luteinizing hormone and follicle-stimulating hormone in the anterior pituitary (Herbison, 2018). Unlike other neurons of the central nervous system, GnRH neurons are born in the olfactory placodes and migrate into the forebrain prenatally (Casoni et al., 2016; Schwanzel-Fukuda et al., 1989; Wray et al., 1989). Recent developmental studies on embryos/fetuses determined the detailed spatio-temporal profile of this process in the human (Casoni et al., 2016). ∼2,000 neurons were observed along a ventral migratory path whereby GnRH neurons reach the hypothalamus to regulate reproduction after puberty. In addition, a previously unknown dorsal migratory route was identified whereby ∼8,000 GnRH neurons migrated towards pallial and/or subpallial structures. The fate of these neurons at later stages of pre- and postnatal development has been unexplored so far.

While GnRH neurons in adult laboratory rodents are mostly hypothalamic and serve reproductive functions (Merchenthaler et al., 1980), a handful of anatomical studies on primates identified additional *GnRH1* mRNA expression and/or GnRH immunoreactivity in extrahypothalamic regions unrelated to reproduction. These included several basal ganglia and the basal forebrain (Krajewski et al., 2003; Quanbeck et al., 1997; Rance et al., 1994; Terasawa et al., 2001). Initial enthusiasm to study these neurons further faded after suggestions that extrahypothalamic GnRH neurons in monkeys contain the GnRH degradation product GnRH1-5, instead of the *bona fide* GnRH decapeptide (Quanbeck et al., 1997; Terasawa et al., 2001).

Here we study human extrahypothalamic GnRH neurons in adult *postmortem* brains with immunohistochemistry (IHC), *in situ* hybridization (ISH), single-cell transcriptomics (RNA-Seq) and high performance liquid chromatography/tandem mass spectrometry (HPLC-MS/MS). We report and characterize a previously unexplored large GnRH neuron population with ∼150,000-200,000 cell bodies scattered in different basal ganglia and the basal forebrain. GnRH neurons of the putamen (Pu) contain *bona fide* GnRH decapeptide, as shown by HPLC-MS/MS. Deep transcriptome analysis reveals that these neurons express GnRH biosynthetic enzymes, *GNRHR1* autoreceptors, inhibitory G proteins implicated in GnRHR1 signaling and the molecular machinery of cholinergic and GABAergic co-transmission. To obtain an insight to the function of GnRH/GnRHR1 signaling in these neurons, we introduce a neonatal mouse model. Slice electrophysiological studies of this model reveal that GnRHR1 autoreceptor activation reduces the resting membrane potential and the electric activity of cholinergic interneurons in the caudate-putamen. Altogether, these data indicate that GnRH is a cotransmitter of many cholinergic neurons in the human Pu and other extrahypothalamic sites. GnRH acts on GnRHR1 autoreceptors to regulate higher order non-reproductive functions associated with cholinergic systems of the basal ganglia and the basal forebrain. Based on these observations GnRH/GnRHR1 signaling may emerge as a new therapeutic target in the treatment of neurodegenerative disorders affecting cholinergic neurocircuitries, including Parkinson’s and Alzheimer’s diseases.

## Results

### Human extrahypothalamic GnRH-immunoreactive neurons occur in the basal ganglia and the basal forebrain

The primate central nervous system contains extrahypothalamic GnRH cell populations which have unknown functions (Krajewski et al., 2003; Quanbeck et al., 1997; Rance et al., 1994; Terasawa et al., 2001). ISH studies of adult human brains identified ∼6-7 thousand *GNRH1* mRNA expressing neurons in the Pu and the nucleus basalis magnocellularis of Meynert (nbM), among other sites (Rance et al., 1994). Here we used IHC to address the presence and map the distribution of GnRH-immunoreactive (IR) neurons in extrahypothalamic sites of adult human brains (N=3). Every 24^th^ 100-µm-thick coronal section between Bregma levels −22.5 and 33.1 (Mai et al., 1997) was immunostained using a well-characterized guinea pig antiserum (#1018) against GnRH decapeptide (Hrabovszky et al., 2011). This experiment revealed numerous extrahypothalamic GnRH-IR neurons in the Pu, moderate numbers in the nucleus accumbens (nAcc) and the head of the nucleus caudatus (Cd) and lower numbers also in the nbM (**Fig. 1A**). Scattered labeled neurons occurred in the globus pallidus (GP), the ventral pallidum (VP) and the bed nucleus of the stria terminalis (BnST). Labeled perikarya showed round or oval shape, with a mean diameter of 29 µm in the Pu (**Fig. 1B**).

**Fig. 1.**
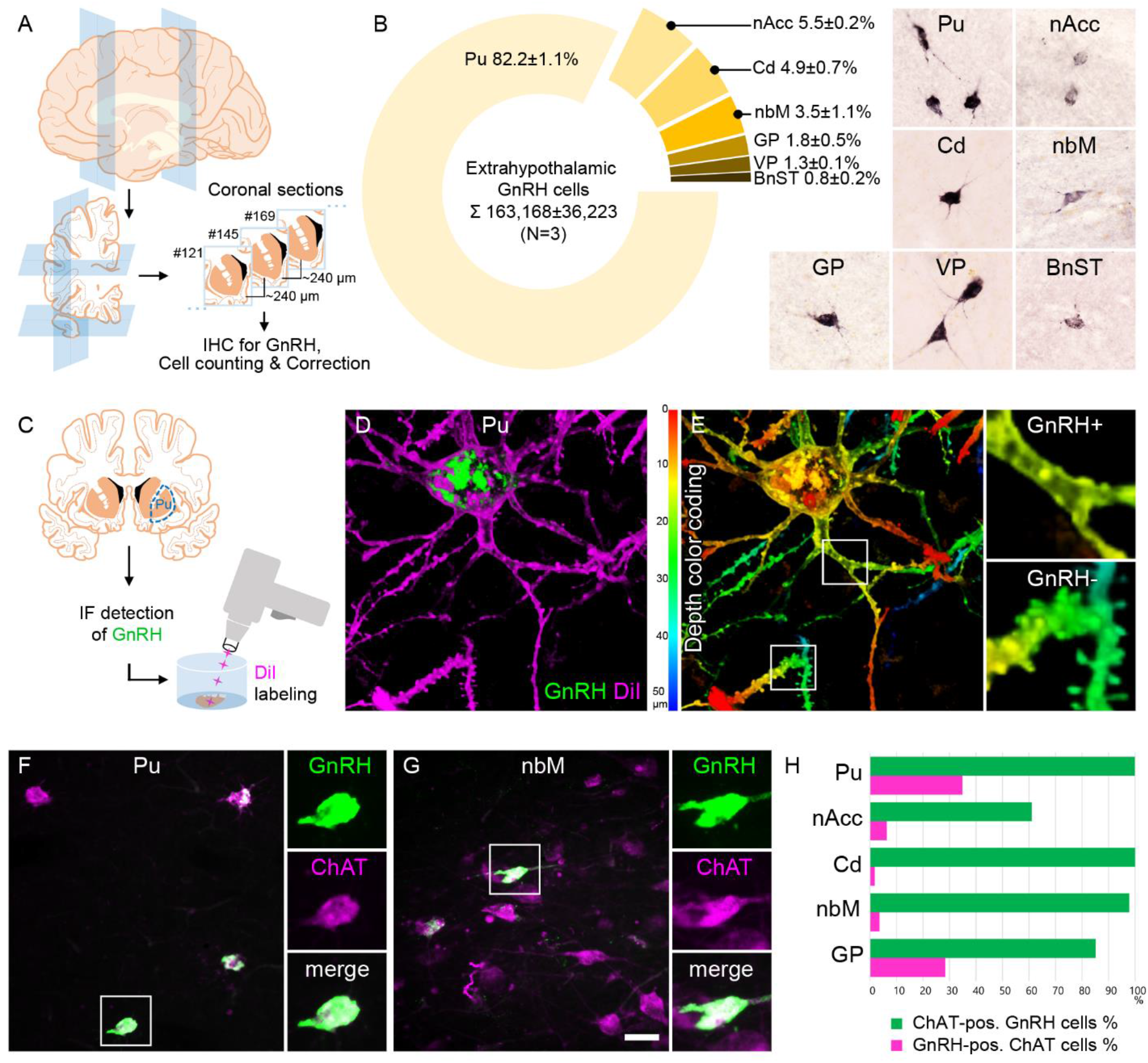
Anatomical approaches unveil the distribution, number, fine structure and cholinergic phenotype of extrahypothalamic GnRH neurons in the adult human brain. **A:** Extrahypothalamic GnRH-IR neurons were mapped with immunohistochemistry (IHC) and quantified in 3 adult human brains. B: The majority (82.2%) of the 163,168±36,223 extrahypothalamic GnRH neurons occurred in the putamen (Pu), followed by the nucleus accumbens (nAcc), nucleus caudatus (Cd), nucleus basalis magnocellularis (nbM), globus pallidus (GP), ventral pallidum (VP) and bed nucleus of the stria terminalis (BnST). **C:** Immunofluorescent (IF) detection of GnRH was combined with cell membrane labeling using Gene Gun-delivered DiI to visualize the fine structure of dendrites. **D:** 3-D reconstruction of the DiI-labeled (magenta) GnRH-immunoreactive (green) neurons revealed large multipolar cells which exhibited only few dendritic spines. **E:** Depth color coding allowed better distinction between DiI-labeled processes of the GnRH neuron (upper inset; GnRH+) from other DiI-labeled neuronal elements many of which belonged to medium spiny GABAergic projection neurons (lower inset; GnRH-). **F:** Double-IF experiments addressed the presence of known interneuron phenotype markers in GnRH neurons. Nearly all GnRH neurons in the Pu contained the cholinergic marker enzyme choline acetyltransferase (ChAT). **G:** The GnRH neuron population also overlapped with cholinergic projection neurons of the nbM. **H:** With few exceptions, GnRH neurons were ChAT-immunoreactive (green columns), whereas they represented smaller subsets of cholinergic cells (magenta columns) being highest in the Pu (∼35%). Scale bar: 50 µm in **B, F, G**, 25 µm in **F, G** insets, 12.5 µm in **D, E** (insets: 3.7 µm).

### Quantitative analysis detects 150-200,000 extrahypothalamic GnRH neurons in the adult human brain most of which are located in the putamen

GnRH neurons develop in the olfactory placodes and migrate to the brain prenatally (Schwanzel-Fukuda et al., 1989; Wray, 2001). Recent studies from Casoni and colleagues identified 10,000 migrating GnRH neurons in human embryos/fetuses most of which (∼8,000) followed a previously unknown dorsal migratory route targeting subpallial and/or pallial structures, instead of the hypothalamus (Casoni et al., 2016). We addressed the possibility that these neurons give rise to extrahypothalamic GnRH-IR neurons of the adult brain by determining the total number of GnRH-IR neurons in the basal ganglia and the basal forebrain. Immunolabeled neurons were counted in every 24^th^ section of a single hemisphere using light microscopy (**Fig. 1A**). Cell counts were then multiplied by 24 and 2 (for two hemispheres) and compensated for overcounting (Abercrombie, 1946; Guillery, 2002) (**Supplementary File 1**). The total number of extrahypothalamic GnRH neurons calculated this way was 229,447, 155,357 and 104,699 in three brains (**Supplementary File 4**), respectively (163,168±36,223; Mean±SEM). Such high cell numbers argued against the placodal origin of extrahypothalamic GnRH neurons. 82.2±1.1 % of labeled cells were observed in the Pu, 5.5±0.2 % in the nAcc, 4.9±0.7 % in the Cd, 3.5±1.1 % in the nbM, 1.8±0.5 % in the GP, 1.3±0.1 % in the VP and 0.8±0.2 % in the BnST (**Fig. 1B**).

### Extrahypothalamic GnRH neurons synthesize *bona fide* GnRH decapeptide derived from the *GNRH1* transcript

Results of previous IHC studies on rhesus monkeys questioned whether extrahypothalamic GnRH neurons synthesize *bona fide* GnRH decapeptide (Quanbeck et al., 1997; Terasawa et al., 2001). First, these cells were not recognized by several GnRH antibodies including the widely-used LR-1 rabbit GnRH polyclonal antiserum (Silverman et al., 1990). Second, they exhibited immunoreactivity to EP24.15 metalloendopeptidase which cleaves GnRH at the Tyr5-Gly6 position to generate GnRH1-5. Here we tested a series of polyclonal antibodies against human GnRH-associated peptide (hGAP1) or GnRH decapeptide (**Supplementary File 5**) for their reactivity with GnRH neurons of the Pu (N=10). All of these antibodies, including the LR-1 antiserum, recognized GnRH-IR neurons (**Supplementary File 2A**), suggesting these cells contain *bona fide* GnRH. Neurons detected with different antibodies were identical as they were double-labeled in dual-immunofluorescence (IF) experiments using two GnRH antibodies from different host species (**Supplementary File 2B**). Results of further experiments with the combined use of IF and non-isotopic ISH showed that GnRH-IR neurons express *GNRH1* mRNA (**Supplementary File 2C**). To provide direct evidence for the biosynthesis of the GnRH decapeptide in these cells, tissue samples were microdissected from the mediobasal hypothalamus (MBH), Pu, Cd and Cl. HPLC-MS/MS analysis of the tissue extracts established that the dominant peptide form in the Pu and Cd is the GnRH decapeptide, with nearly 4-times lower tissue concentrations of GnRH1-5. Only GnRH decapeptide was detectable in the MBH (used as a positive control) where hypophysiotropic GnRH neurons occur and neither peptide form was present in the Cl, in accordance with the absence of IHC labeling at this site (**Supplementary File 2D-F**). Together with observations from the IHC and ISH experiments, HPLC-MS/MS results gave firm support to the notion that extrahypothalamic GnRH neurons synthesize *bona fide* GnRH decapeptide derived from the *GNRH1* gene.

### GnRH neurons of the putamen are large multipolar interneurons with smooth-surfaced dendrites

The immunohistochemical method left important fine structural properties of extrahypothalamic GnRH-IR cells unlabeled (**Fig. 1B**). Therefore, the dendritic compartment was labeled with the lipophilic dye DiI for further analysis (Takacs et al., 2018) (**Fig. 1C**). Following the immunofluorescent visualization of GnRH neurons in the Pu of a 72-year-old female, DiI-coated tungsten particles were delivered into the sections using a Helios Gene Gun (Bio-Rad) (Takacs et al., 2018). Spreading of this lipophilic dye along the cytoplasmic membrane surface caused Golgi-like labeling of random-hit neurons, including 12 GnRH-IR cells (**Fig. 1D**). Confocal microscopic analysis and 3-D reconstruction of the DiI signal revealed spider-like neurons with rich arborization of poorly-spined dendrites. DiI-labeled GnRH neurons were clearly distinct from the main Pu cell type, the densely-spined medium spiny GABAergic projection neuron (SPN) (**Fig. 1E**).

### Extrahypothalamic GnRH cells represent subpopulations of cholinergic neurons

SPNs represent 80-98% of striatal neurons, the remainder being made up of cholinergic and different subclasses of GABAergic interneurons (Gonzales et al., 2015). DiI-labeled GnRH cells resembled anatomically to cholinergic interneurons (ChINs). Indeed, dual-IF experiments established that GnRH neurons of the Pu contain the cholinergic marker enzyme choline acetyltransferase (ChAT) (**Fig. 1F**). Similarly, GnRH neurons in the nbM (**Fig. 1G**) and other extrahypothalamic sites contained ChAT immunoreactivity. The extent of ChAT/GnRH colocalization was assessed quantitatively in five distinct regions of a 62-year-old male subject (#3). Confocal microscopic analysis of representative dual-labeled sections established that the vast majority of extrahypothalamic GnRH neurons are cholinergic (**green bars in Fig. 1H**). In contrast, GnRH-IR neurons represented only 34.9% of all cholinergic neurons in the Pu, 1.8% in the head of the Cd, 6.3% in the nAcc, 28.4% in the GP and 3.6% in the nbM (**magenta bars in Fig. 1H**). GnRH-positive and GnRH-negative cholinergic neurons often intermingled, without gross morphological differences between the two phenotypes (**Figs. 1F, G**).

### Hypothalamic GnRH neurons regulating reproduction also exhibit an unexpected cholinergic phenotype

The ChAT phenotype emerged as a hallmark of extrahypothalamic GnRH neurons. To verify absence of ChAT in the hypothalamic GnRH neuron population, tissue sections were processed for dual-IF detection of the ChAT and GnRH antigens and analyzed with confocal microscopy. Unexpectedly, 41.2±7.1% of the hypothalamic GnRH neurons also exhibited ChAT signal in adult human male and female subjects (N=7) (**Fig. 2A**), a phenomenon not observed in other species before.

**Fig. 2.**
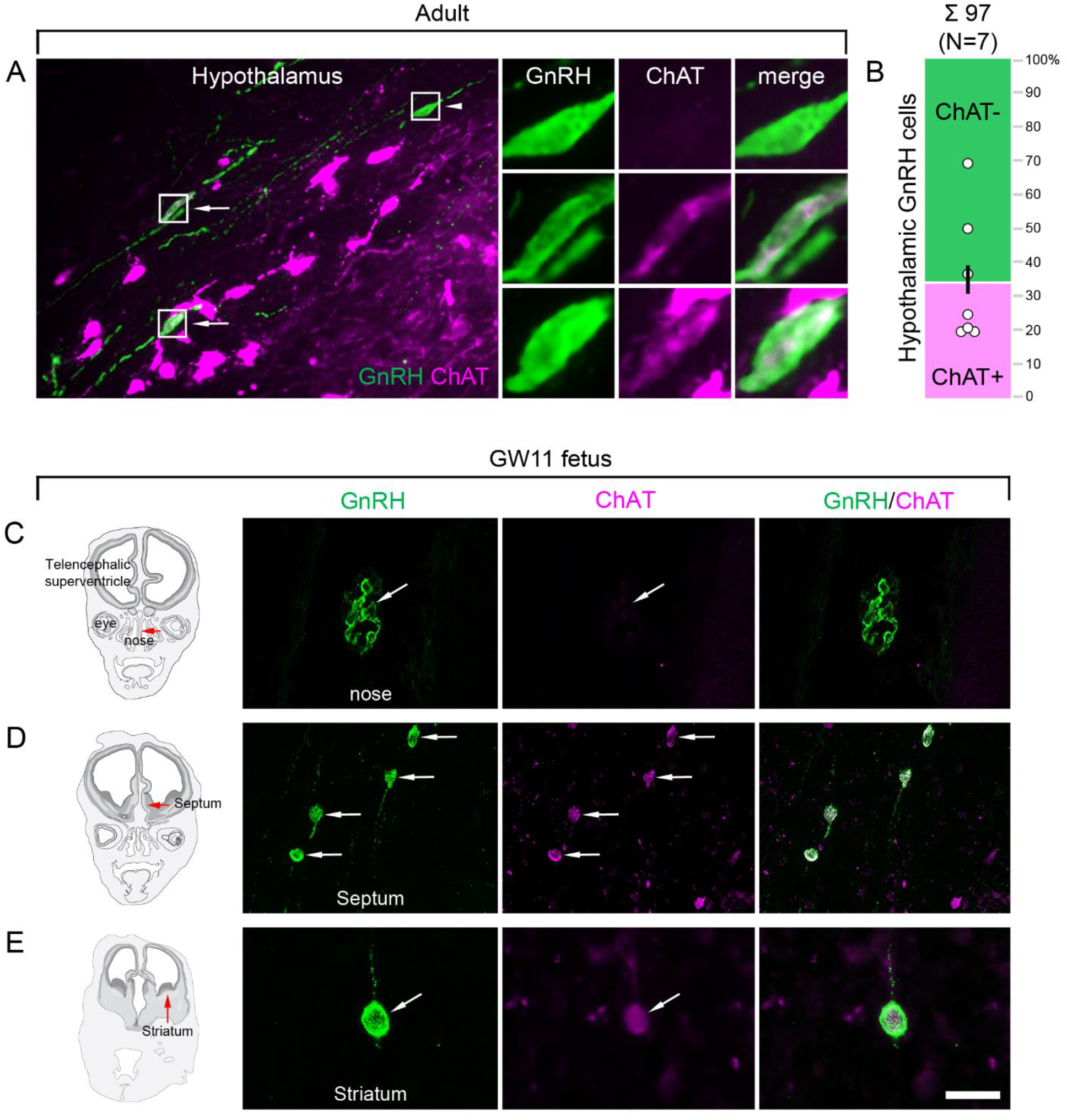
Both hypothalamic and extrahypothalamic GnRH neurons exhibit cholinergic phenotype gained during early fetal development. **A:** The cholinergic phenotype is not a hallmark of extrahypothalamic GnRH neurons because large subsets of GnRH neurons in the adult human hypothalamus (green immunofluorescent signal) also exhibit choline acetyltransferase (ChAT; magenta) immunoreactivity. High-power insets show single-(arrowhead) and dual-labeled (arrows) GnRH neurons from framed regions. **B:** Quantitative analysis of 97 GnRH neurons from 7 subjects reveal the ChAT phenotype in 41.2±7.1% of hypothalamic GnRH neurons. **C-E:** The cholinergic phenotype of GnRH neurons is gained during early fetal development. Left panels illustrate coronal views of the fetal head at GW11. Representative photomicrographs taken from sites indicated by the red arrows show results of dual-immunofluorescence experiments. **C:** At this stage of development a large subset of GnRH neurons (green immunofluorescent signal) migrate in the nasal region toward the brain and do not exhibit ChAT signal. **D, E:** In contrast, GnRH neurons migrating through the septal area (**D**, arrows) or located in the striatum (**E**, arrow) express ChAT (magenta). Scale bar: 50 µm in **A** (insets: 12.5 µm), **C** and **D**, 20 µm in **E**.

### Cholinergic phenotype of GnRH neurons develops prenatally

Prenatal co-expression of ChAT and GnRH was then explored via dual-IF experiments in coronal sections of fetal heads (N=2) at gestational week 11 (GW11). At this age ∼20% of GnRH neurons can still be found in the nasal region, whereas the majority have already entered the brain to migrate toward hypothalamic and extrahypothalamic target areas (Casoni et al., 2016). While GnRH positive neurons within the nasal compartment did not contain ChAT signal (**Fig. 2B**), those in the septum (**Fig. 2C**), the striatum (**Fig. 2D**) and elsewhere in the developing brain were already ChAT-IR. These data suggest that migrating GnRH neurons gain their cholinergic phenotype as soon as they enter the brain and continue to express ChAT immunoreactivity in hypothalamic as well as extra-hypothalamic regions.

### Transient GnRH-GFP transgene expression in the caudate-putamen of neonatal mice offers a functional model

Functional studies of extrahypothalamic GnRH neurons require relevant animal models. Although GnRH immunoreactivity or mRNA expression has never been reported in the rodent caudate-putamen (CPU), in a pilot study we noticed that the developing CPU of a GnRH-enhanced green fluorescent protein (GnRH-GFP) transgenic mouse strain (Suter et al., 2000) transiently expresses green fluorescence. The fluorescent signal was most intense at postnatal week 1 (PNW1) and then, gradually faded to disappear by PNW4 (**Supplementary File 3A-C**). ChAT immunoreactivity showed an inverse temporal profile, being nearly undetectable at PNW1 and increasing with time (**Supplementary File 3A-C**). As established in PNW2 mice, the transient GnRH-GFP fluorescence characterized selectively a subset of the ChINs in the CPU (**Supplementary File 3B**). Although efforts to confirm neonatal GnRH biosynthesis with HPLC-MS/MS or RT-PCR in the PNW1 CPU failed likely because of the low mRNA and peptide expression levels, the transient GnRH-GFP transgene expression of ChINs raised the possibility that neonatal mice are a relevant model to study GnRH effects in the striatum via slice electrophysiology. Three transgenic mouse strains showing green fluorescence selectively in GnRH-GFP neurons, in cholinergic neurons (ChAT-Cre/zsGreen) and in GABAegic neurons (GAD65-GFP) (Lopez-Bendito et al., 2004), respectively, were used.

### GnRH inhibits GnRH-GFP and ChAT-Cre/zsGreen neurons in the CPU of neonatal mice

In whole-cell patch-clamp experiments on PNW1 mice (**Fig. 3A**), 7 out of 15 GnRH-GFP neurons responded to 1.2 µM GnRH with a transient hyperpolarization (V_rest_ = −51.0 ± 1.11 mV, ΔV_rest_ = −4.3 ± 0.99 mV, **Fig. 3B**; p=0.0007) which started within 2.7 ± 2.1 min and persisted for 8.0 ± 4.5 min. The majority of ChINs were silent at resting potential. Therefore, action potentials (APs) were induced by a 10 pA/15-min-long depolarizing current pulse to address GnRH effects on neuronal activity. GnRH transiently decreased the firing rate in 7 out of 13 GnRH-GFP neurons to 69.3 ± 10.01% of the control value (1.36 ± 0.06 Hz, **Fig. 3B**; p=0.0098). GnRH elicited similar inhibitory responses from ChINs of PNW1 ChAT-Cre/zsGreen mice and hyperpolarized 8 out of 15 cholinergic neurons (V_rest_ = −53.6 ± 2.48 mV, ΔV_rest_ = −4.1 ± 0.87 mV, **Fig. 3C**; p=0.0004). Furthermore, in 7 out of 13 neurons, GnRH decreased transiently the current pulse-induced firing activity to 72.5 ± 7.61% of the 1.05 ± 0.13 Hz control value (**Fig. 3C**; p=0.0098).

**Fig. 3.**
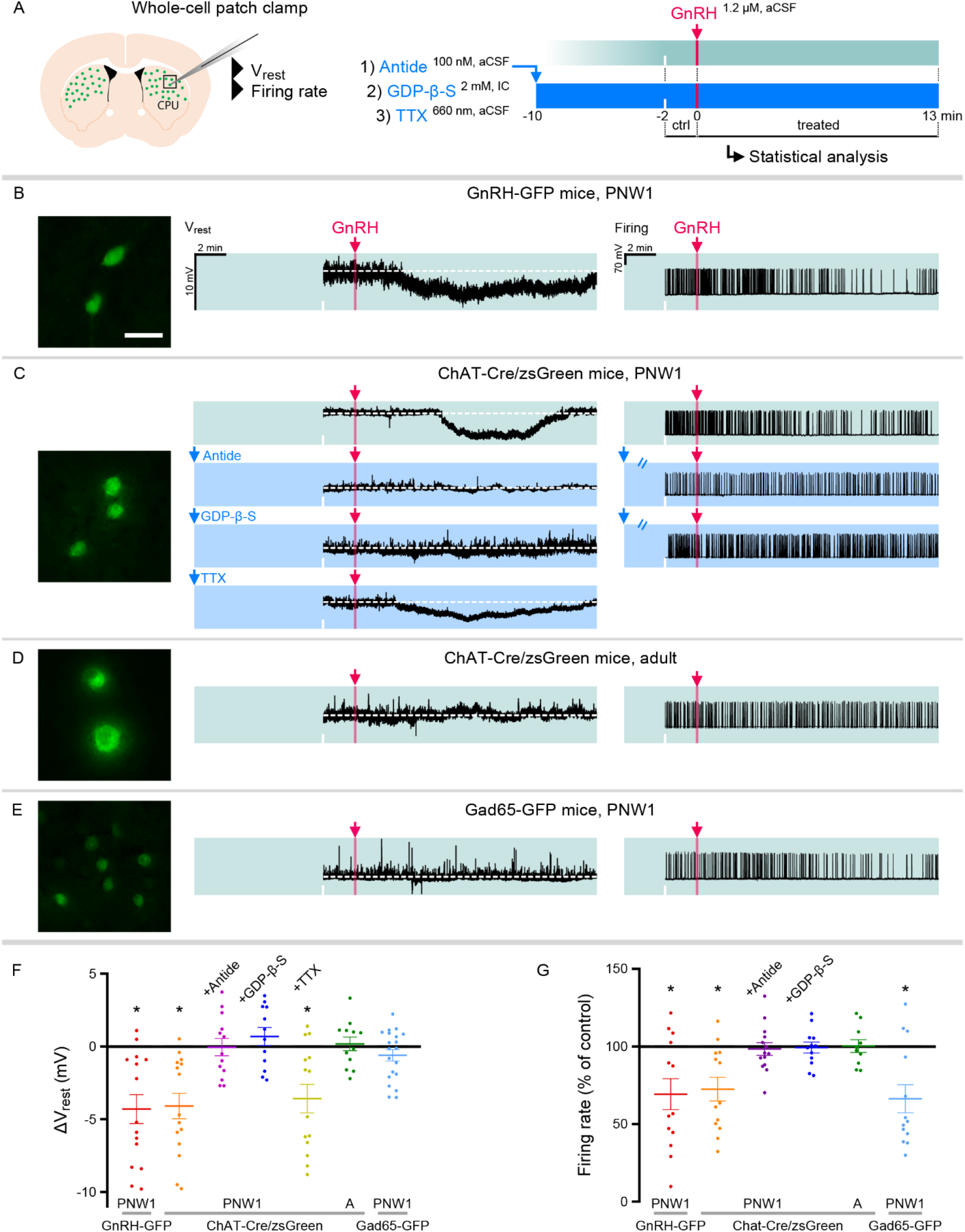
GnRH inhibits cholinergic interneurons in the caudate-putamen of neonatal transgenic mice via signaling on GnRHR1 autoreceptors. **A:** Neonatal GnRH-GFP transgene expression within cholinergic neurons of the caudate-putamen (CPU) indicates that newborn transgenic mice may serve as animal model to study GnRH effects with slice electrophysiology. To reveal receptor mechanisms, the selective GnRHR1 antagonist Antide, the membrane-impermeable G Protein-Coupled Receptor inhibitor GDP-β-S and the action potantial inhibitor tetrodotoxin (TTX) were used in whole-cell patch-clamp experiments. **B:** Postnatal week 1 (PNW1; postnatal day 4-7) GnRH-GFP neurons responded to GnRH with reduced resting membrane potential (V_rest_) and decreased rates in current pulse-induced firing activity. **C:** The same inhibitory responses could also be elicited from cholinergic interneurons of newborn ChAT-Cre/zsGreen. GnRH acted via its specific receptor GnRHR1 because inhibitory responses could be prevented with Antide. GnRHR1 mediating GnRH effects was localized within CPU cholinergic neurons. First, GnRH was unable to inhibit cholinergic neurons if the internal electrode solution contained GDP-β-S. Second, GnRH was still able to hyperpolarize cholinergic neurons in the presence of TTX to eliminate activity-dependent indirect actions (TTX+GnRH). **D:** In contrast to the newborn mice, adult ChAT-Cre/zsGreen animals did not respond to GnRH with reduced V_rest_ or firing rate. **E:** Medium spiny projection neurons which receive input from cholinergic interneurons were studied in neonatal GAD65-GFP transgenic mice. GnRH did not change the V_rest_ but decreased the firing rate of these neurons, indicating together that the inhibitory response is indirect. **F, G:** Scatter dot plots summarize the results of measurements in the different treatment groups. *p<0.05 with ANOVA. Scale bar: 25 µm. See also **Figure 3 – Source Data** for recordings.

### GnRH-dependent inhibition of neonatal cholinergic neurons is mediated by GnRH receptors

To identify the receptor involved in the GnRH-induced inhibition of neonatal ChAT-Cre/zsGreen neurons, slices were preincubated with the selective GnRH receptor (GnRHR1) antagonist Antide (100 nM) for 10 min prior to GnRH administration (**Fig. 3A**). In the presence of Antide, GnRH was unable to affect the resting membrane potential (N=13; **Fig. 3C**) and the firing rate (N=14; **Fig. 3C**) of ChINs, indicating that GnRH acts on GnRHR1. Neurons containing GnRHR1 remained to be identified.

### GnRHR1 is localized to ChINs

GnRHR1 is a G-protein coupled receptor (GPCR). When the membrane-impermeable GPCR inhibitor GDP-β-S (2 mM, **Fig. 3A**) was added to the internal electrode solution, GnRH was unable to alter the V_rest_ (N=12) and the firing rate (N=12) of ChINs (**Fig. 3C**). In addition, when the action potential inhibitor TTX (660 nM, **Fig. 3A**) was present in the aCSF to eliminate activity-dependent transsynaptic effects, GnRH was still able to hyperpolarize 7 out of 14 ChINs (V_rest_ = −54.1 ± 1.59 mV, ΔV_rest_ = −3.6 ± 0.97 mV, **Fig. 3C**; p=0.0029). Together, these functional results served as proof that GnRHR1 mediating the effects of exogenous GnRH is localized within ∼50 % of CPU ChINs.

### GnRH does not influence ChINs in adult mice

GnRH actions on ChINs were only observed in PNW1 mice and none of the ChAT-Cre/zsGreen neurons responded with altered V_rest_ (N=12) or firing rate (N=10) to GnRH in adult animals (**Fig. 3D**).

### GnRH inhibits CPU GABAergic neurons via indirect actions

The majority of striatal neurons are medium-sized GABAergic SPNs which receive strong input from ChINs (Gonzales et al., 2015). GnRH did not change the resting membrane potential of putative SPNs (medium-sized GAD65-GFP neurons) in neonatal transgenic mice (N=20; **Fig. 3E**). In turn, GnRH decreased the firing rate of 9 from 13 GAD65-GFP neurons to 66.3 ± 9.07% of the control frequency (0.84 ± 0.16 Hz, **Fig. 3E**; p=0.0030). Together, these observations suggested that GnRH can inhibit a subset of SPNs via acting indirectly.

ANOVA revealed significant effects of GnRH on V_rest_ and firing rates in neonatal GnRH-GFP and ChAT-Cre/zsGreen neurons but not in adult ChAT-Cre/zsGreen neurons (**Figs. 3F, G**). Application of Antide or GDP-β-S alone did not change the V_rest_ or the firing rate of the control recording periods (see **Figure 3 – Source Data**).

### Neurons laser-capture microdissected from the *postmortem* putamen provide sources for high-quality RNA suitable for RNA-seq

Electrophysiological observations indicated that GnRH acts on inhibitory GnRHR1 autoreceptors within CPU ChINs of newborn mice. To address if a similar GnRHR1 autoreceptor signaling also acts in the adult human Pu, deep transcriptome profiling of ChINs was carried out using SPNs as a control cell population. Being the largest cell type, ChINs were readily recognizable in sections subjected to Nissl-staining under RNase-free conditions (**Fig. 4A**). Laser capture microdissection (LCM) was used to collect neurons from cresyl violet-stained Pu sections of two human subjects. Each ChIN-enriched pool contained ∼300 large neurons a third of which are GnRH-IR (**Fig. 1H**). Each of the two SPN-enriched control pools consisted of ∼600 medium-sized neurons (**Fig. 4A**). Total RNA was isolated and RNA-Seq libraries prepared from the four cell pools and sequenced with the Illumina NextSeq 500/550 High Output (v2.5) kit. 23.4 M and 18.4 M reads were obtained from the two ChIN pools, respectively, from which ∼9.6 M and 6.6 M reads were mapped to transcripts of the the GRCh38.p13 human reference genome; 13664 and 12637 transcripts occurred at cpm >5 in ChINs (**Fig. 4A**).

**Fig. 4.**
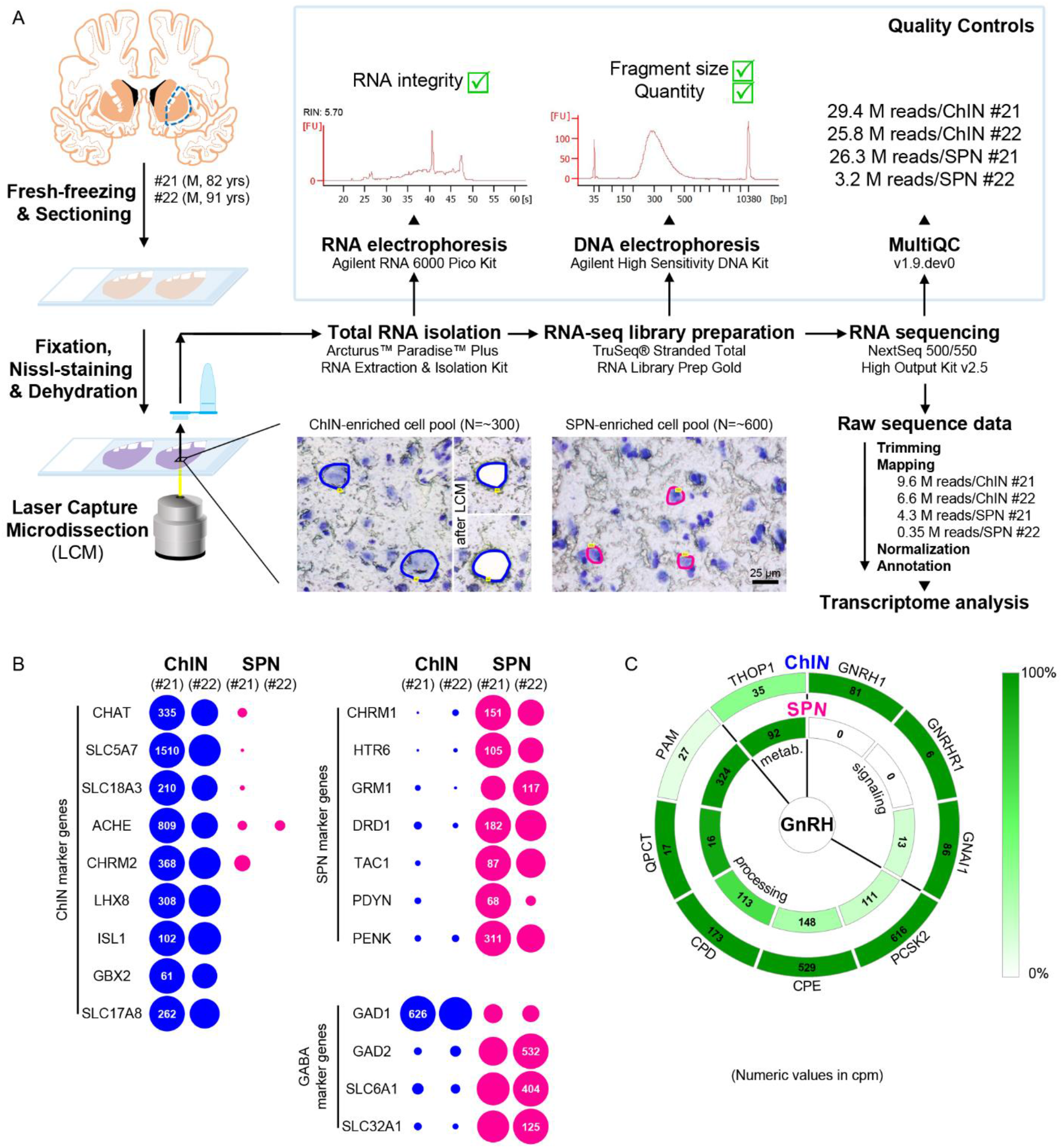
Deep transcriptome profiling of cholinergic interneurons and spiny projection neurons provides new insight into extrahypothalamic GnRH signaling mechanisms and the molecular connectome of the human putamen. **A:** 20-µm-thick coronal sections were collected on PEN membrane slides from frozen putamen samples of two male human subjects (#21 and 22) and fixed with an ethanol/paraformaldehyde mixture. Neurons were visualized using Nissl-staining and isolated with laser-capture microdissection (LCM). 300 neurons included in each cholinergic interneuron (ChIN)-enriched cell pool were recognized based on their large perikaryon size. 600 medium-sized neurons in the control pool mostly included medium spiny GABAergic projection neurons (SPNs), the major putamen cell type. Total RNA was isolated and RNA-Seq library prepared from both cell pools and sequenced with the Illumina NextSeq 500/550 High Output (v2.5) kit. **B:** Bioinformatic analysis verified high enrichment of known cholinergic markers in the two ChIN pools and of SPN markers in the SPN control pools. Expression levels in dots reflect counts per million reads (cpm) and in each case, dot areas reflect transcript abundances relative to the highest cpm (100%). **C:** Key elements of proGnRH processing, GnRH signaling and GnRH metabolism are illustrated in two concentric circles. The *GNRH1* and *GNRHR1* transcripts are present in ChINs only (outer circle). ChINs express inhibitory G protein-coupled receptor subnunits including *GNAI1*, all enzymes required for proGnRH processing and *THOP1* which may account for GnRH cleavage. Color coding reflects relative transcript abundances, whereas numbers indicate cpms (mean cpms of subjects #21 and 22).

### Size-based laser-capture microdissection allows adequate sampling of striatal cholinergic interneurons and medium spiny projection neurons

Cholinergic markers, including *CHAT*, *SLC5A7*, *SLC18A3*, *ACHE* and *CHRM2*, were highly enriched in the ChIN pools from subjects #21 and #22. These transcripts were either absent or found at low levels only in the two SPN pools (**Fig. 4B**). Mouse ChINs arise from Nkx2.1+ progenitors. During development, Nkx2.1 upregulates the expression of the LIM homeobox proteins LHX8, ISL1 and GBX2 which, in turn, promote cell differentiation into ChINs (Allaway et al., 2017). These LIM transcripts as well as type-3 vesicular glutamate transporter (*SLC17A8*) showed robust and exclusive expression in ChINs (**Fig. 4B**). The control SPN pools, in turn, expressed much higher levels of known SPN markers than ChINs, including cholinergic (*CHRM1*), serotonergic (*HTR6*), glutamatergic (*GRM1*) and dopaminergic (*DRD1*) receptor isoforms and several neuropeptides (*TAC1*, *PDYN*, *PENK*) (**Fig. 4B**). Differential distribution of the above transcripts verified that the size-based LCM strategy efficiently separated ChINs from SPNs for deep transcriptome profiling. Relatively high levels of expression of known GABAergic marker transcripts *GAD1*, *GAD2*, *SLC32A1* and *SLC6A1* in ChINs, in addition to SPNs (**Fig. 4B**), revealed that ChINs use GABAergic co-transmission, as proposed earlier in the rodent CPU (Lozovaya et al., 2018).

### Cholinergic interneurons selectively express *GNRH1* and *GNRHR1* and contain GnRH biosynthetic enzymes and inhibitory G proteins

*GNRH1* was expressed exclusively in the two ChIN pools, confirming the morphological observations (**Fig. 4C**). Processing of the proGnRH1 protein begins with endoproteolysis by prohormone convertases from which ChINs abundantly expressed the *PCSK2* isoform. Enzymes catalyzing subsequent steps of GnRH biosythesis, including carboxypeptidases (*CPE*, *CPD*), peptidylglycine α-amidating monooxygenase (*PAM*), and glutaminyl cyclase enzymes (*QPCT*), were also present in ChINs (**Fig. 4C**). The THOP1 peptidase accounts for the degradation of multiple neuropeptides, including GnRH. This enzyme was expressed in both ChINs and SPNs, with a higher abundance in the latter. The seven transmembrane receptor *GNRHR1* was expressed selectively in ChINs, strongly suggesting that GnRH in the human Pu acts on GnRHR1 autoreceptors. Electrophysiological observations on newborn mice showed that this action is mediated by inhibitory G-proteins encoded by *GNAI* genes. Indeed, GnRHR1 can be coupled to inhibitory G-proteins in prostate cancer (Limonta et al., 1999) and in GT1-7 cells (Krsmanovic et al., 2003). We found that ChINs expressed all three *GNAI* isoforms, with the highest abundance of *GNAI1* (**Fig. 4C**). Altogether, transcriptome profiling of ChINs and SPNs provided molecular support to the concept that GnRH is synthesized by ChINs and acts via inhibitory G-protein-coupled GnRHR1 autoreceptors.

### Transcriptome profiling provides novel insight into the molecular connectome of the human putamen

Deep transcriptome profiling of ChINs and SPNs revealed a large set of genes that were expressed selectively or predominantly in one cell type only, in addition to many other genes expressed in both. Neurotransmitter and neurotransmitter receptor transcripts identified this way allowed us to propose signaling mechanisms that act in the bidirectional communication between ChINs and SPNs. Some receptors appear to serve as autoreceptors (e.g. GnRHR1, NMBR, CRH1R/2R). Others may receive ligands from multiple neuronal sources within (e.g.: QRFPR, NPY1R/5R, TACR1, SSTR2/3) or outside (e.g.: OXTR, MC4R, GLP1R, PRLR1) the striatum. Peptidergic mechanisms concluded from the transcriptome profiles are illustrated as a schematic model in **Fig. 5**. A deeper insight into the molecular connectome of the human Pu can be obtained from the detailed receptor and neuropeptide expression profiles of ChINs and SPNs (**Supplementary File 6**; BioProject accession number: PRJNA680536)

**Fig. 5.**
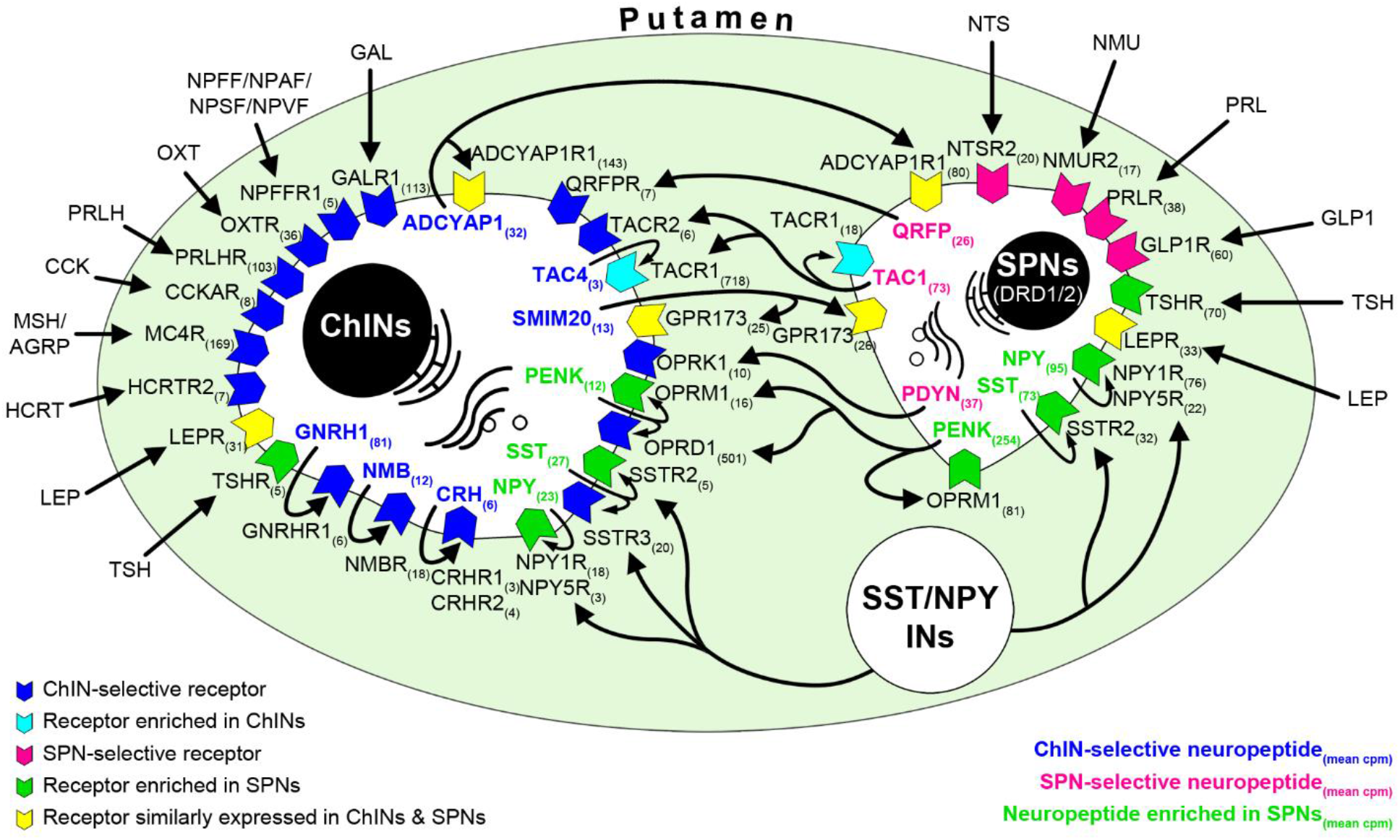
RNA-Seq studies reveal the neuropeptide and neuropeptide receptor expression profiles of cholinergic interneurons and spiny projection neurons and provide insight into the molecular connectome of putamen cell types. Proposed signaling mechanisms are based on neuropeptide and peptide receptor expression profiles of the two cell types. ChINs appear to use GnRHR1, CRH1R/2R and NMBR autoreceptor signaling. SSTR2, NPY1R/5R, NKR1, OPRM1 and TACR1 may serve, at least partly, as autoreceptors in SPNs. Proposed peptidergic communication between the two cell types are also indicated by arrows. Other receptors receive ligands from different neuronal sources within (e.g.: QRFPR, NPY1R/5R, TACR1, SSTR2/3) or outside (e.g.: OXTR, MC4R, GLP1R, PRLR1) the putamen. Numbers in receptor symbols reflect transcript abundances expressed as mean CPMs (counts per million) from subjects #21 and #22. The figure illustrates receptors that were consistently observed in the given cell type of both human samples. Abbreviations: ChINs, cholinergic interneurons; INs, interneurons; SPNs, spiny projection neurons.

## Discussion

### Extrahypothalamic GnRH-IR neurons correspond to Type-III GnRH neurons detected earlier with *in situ* hybridization

A pioneer ISH study by Rance and co-workers distinguished three types of *GNRH1* mRNA expressing neurons in the human brain based on size, shape, and labeling intensity (Rance et al., 1994). GnRH-IR neurons detected in our study correspond to Type-III neurons characterized by round/oval shape, large nucleus and nucleolus, prominent Nissl substance and *GNRH1* mRNA levels intermediate between those of heavily-labeled Type-I neurons in the mediobasal hypothalamus and lightly-labeled Type-II neurons in the medial septum and the dorsal medial preoptic area (Rance et al., 1994). Our IHC also detected many heavily labeled Type-I hypothalamic GnRH neurons but only few septal Type-II neurons which latter had negligeable contribution to the total GnRH cell numbers. Type-III neurons also occur in non-human primates (Krajewski et al., 2003; Quanbeck et al., 1997; Terasawa et al., 2001), whereas they have not been reported in rodent species.

### Overlap with cholinergic neurons and large cell numbers argue against the placodal origin

The ISH study of Rance and coworkers identified 5,800 Type-III GnRH neurons in the basal forebrain complex rostral to the mammillary bodies, caudal to the optic chiasm and ventral to the anterior commissure (Rance et al., 1994). Tissues with these anatomical guidelines are devoid of the bulk of the Pu which contained the majority (82%) of the extrahypothalamic GnRH neurons in our study. Total GnRH-IR cell numbers we calculated for the basal forebrain and basal ganglia of three adult brains (229,447, 155,357 and 104,699, respectively) exceeded all previous estimates and also made it unlikely that these cells are identical to GnRH neurons observed recently along the dorsal migratory route (∼8,000) during embryonic/fetal development (Casoni et al., 2016). Extrahypothalamic GnRH neurons of the human seem to be homologous to the early type of developing GnRH neurons reported from monkeys (Quanbeck et al., 1997). In this species, the early and late types of GnRH neurons were distinguished based on differences in their time of appearance, morphology and immunoreactivity pattern using GnRH antibodies against different GnRH epitopes (Quanbeck et al., 1997). It was speculated that early GnRH neurons originated from the dorsal olfactory placode before olfactory pit formation at E30, migrated into the brain along the olfactory nerve and settled in striatal and limbic structures of the fetal brain (Quanbeck et al., 1997). However, in a subsequent study (Terasawa et al., 2001) these authors noted that a 10- to 10,000-fold increase in the number of “early” GnRH neurons in the basal forebrain during the next 4 days indicates that early GnRH neurons might be derived from the ventricular wall of the telencephalic vesicle. The possibility of non-placodal GnRH neuron development is compatible with the *in vitro* capability of hypothalamic and hippocampal progenitors to generate GnRH cells and all other neuroendocrine cell types (Markakis et al., 2004).

It is worth noting that our RNA-Seq studies provided transcriptomic information about a mixed ChIN population of the Pu, whereas ChINs exhibit substantial diversity in their physiology, morphology, and connectivity (Gonzales et al., 2015). Subclasses differ in their developmental origin (medial ganglionic eminence, septal epithelium or preoptic area) and transcription factor profiles (Ahmed et al., 2019). It remains to be determined which ChIN subset expresses *GNRH1*. Selective harvesting of intact cellular RNA specifically from GnRH-IR ChINs of the *postmortem* Pu remains a technical challenge.

### Extrahypothalamic GnRH neurons contain the full-length GnRH decapeptide derived from the *GNRH1* gene

It was proposed that extrahypothalamic GnRH neurons of the monkey contain the GnRH1-5 degradation product of GnRH, instead of the *bona fide* GnRH decapeptide (Quanbeck et al., 1997; Terasawa et al., 2001). Circumstantial evidence to support this notion stemmed from the observations that i) these neurons can not be immunolabeled with the LR-1 rabbit polyclonal antiserum and some other antibodies, and ii) they are IR to the THOP1 enzyme which can cleave GnRH at the Tyr5-Gly6 position. In contrast, our results suggest that the human Pu mostly synthesizes *bona fide* GnRH decapeptide. First, its ChINs can be immunolabeled with the LR-1 antibodies (and several other GAP1 and GnRH antibodies). Second, ChINs possess the full enzyme set of GnRH biosynthesis, as revealed by deep transcriptome profiling. Finally, ChINs contain 4-times as much uncleaved GnRH decapeptide as GnRH1-5, as shown by results of HPLC-MS/MS studies. It is worth to note that the human genome contains a fully functional *GNRH2* gene (Stewart et al., 2009), in addition to *GNRH1*. Nevertheless, the GnRH signal we detected in the Pu is due to *GNRH1,* rather than *GNRH2* expression, because i) extrahypothalamic GnRH-IR neurons exhibit ISH signal for *GNRH1* mRNA, ii) they are IR to GAP1 which has low homology with the corresponding GAP2 sequence, iii) and ChINs of the Pu express high levels of *GNRH1*, but not *GNRH2* mRNA, according to RNA-Seq results.

### Both hypothalamic and extrahypothalamic GnRH neurons use cholinergic co-transmission

ChAT co-expression provided evidence that extrahypothalamic GnRH neurons correspond to subpopulations of previously known cholinergic cells. These include ChINs of the Pu which communicate locally with SPNs as well as projection neurons of the nbM which innervate distant limbic structures (Ahmed et al., 2019). Although ChAT emerged as a hallmark of the extrahypothalamic GnRH system, we found evidence that a relatively large subset of human hypothalamic GnRH neurons also express this cholinergic marker enzyme. To our knowledge, this colocalization has not been reported in any other species before, suggesting a species difference from rodent GnRH neurons which are regulated by cholinergic afferents but not known to co-express cholinergic markers (Turi et al., 2008). Our colocalization experiments on GW11 human fetuses established that migratory GnRH neurons in the nasal compartment are not cholinergic, whereas both hypothalamic and extrahypothamic GnRH neurons already express the ChAT signal at this age.

### Neonatal mice may provide tools for functional studies of striatal GnRH signaling

The transient GFP fluorescence we observed in the CPU of neonatal GnRH-GFP transgenic mice (Suter et al., 2000) has not been reported before. Although so far we were unable to confirm GnRH and GnRHR1 biosynthesis in these cells, the identification of the GnRH promoter-driven selective GFP signal in ChINs raised the possibility to use neonatal mice as relevant models to study extrahypothalamic GnRH signaling with electrophysiology. In whole-cell patch-clamp experiments on tissue slices of newborn mice, exogenous GnRH effectively decreased the resting membrane potential and firing activity of a subpopulation (∼50%) of CPU cholinergic cells. The inhibitory action of GnRH was exerted on cholinergic neurons via GnRHR1 autoreceptors, because i) it was prevented by the GnRHR1 antagonist Antide, ii) as well as by the intracellularly applied universal G-protein coupled receptor inhibitor. RNA-Seq detection of selective *GNRHR1* mRNA expression in ChINs of the human Pu suggests that these functional data are relevant to the human. However, we have to recognize that the neonatal mouse model has severe limitations. First, the electrophysiological responses of immature murine CPU neurons are not necessarily relevant to those of adult human ChINs. Second, while the GnRH signal is readily detectable in the adult human Pu using anatomical approaches, no one has so far been able to detect GnRH decapeptide and/or mRNA signals in the CPU of neonatal mice, which likely reflects extremely low levels of expression.

### Laser-microdissection of size-selected cholinergic interneurons and spiny projection neurons is a highly efficient approach to characterize these cell types from the *postmortem* brain

Deep transcriptome profiling of *postmortem* human neurons is technically challenging. Difficulties include i) compromized RNA quality, ii) lack of obvious marker signals to distinguish cell types, and iii) low RNA yield from the LCM-isolated 300-600 neurons. Our strategy to isolate size-selected ChINs and SPNs with LCM was justified by the RNA-Seq results which showed high enrichment of known cell type-specific marker genes in the two cell pools and millions of identified reads in each. As one-third of ChINs in the Pu also contain GnRH, deep transcriptome profiling of ChINs offered an insight into the extrahypothalamic GnRH neuron transcriptome. It is important to recognize that ChINs of the Pu consist of several subclasses (Ahmed et al., 2019; Gonzales et al., 2015) and our RNA-Seq studies determined the transcriptome profile of a mixed ChIN cell population. Thus, it remains to be determined which ChIN subset expresses *GNRH1* and *GNRHR1*. To answer this question, a technical challenge for the future will be to collect intact RNA from GnRH neurons identified immunohistochemically or with ISH.

### The transcriptome profile of cholinergic interneurons and spiny projection neurons provides novel insight into the molecular connectome of the human putamen

Although it was beyond the focus of our study, deep sequencing of ChINs and SPNs also unveiled the neurotransmitter and receptor profiles of these cell types and provided information about the putative molecular interactions taking place in the Pu. The transcriptome databases allowed us to propose putative peptidergic mechanisms and thus, build the partial molecular connectome model of the two cell types.

### GnRH acts outside the hypothalamus to regulate various reproductive and non-reproductive functions

Clearly, the functions of GnRH are far from being restricted to the regulation of hypophysial gonadotropin secretion. Its receptor, *GNRHR1* is expressed in normal peripheral endocrine tissues including the uterus, the placenta, the ovaries, the testes and the prostate gland as well as in various tumour cell types (Harrison et al., 2004). High levels of *GNRHR1* mRNA and immunoreactivity were reported in pyramidal neurons of the human hippocampus and cerebral cortex (Wilson et al., 2006). GnRH analogues were anti-apoptotic in a rat model of ischemia/reperfusion (Chu et al., 2010). Further, GnRH increased hippocampal estradiol levels and the spontaneous firing and *GNRHR1* expression of pyramidal neurons and prevented memory deficits caused by amyloid β deposition (Marbouti et al., 2020). While the source of GnRH acting on hippocampal neurons remains to be explored, GnRHR1 in ChINs of the basal ganglia can bind locally synthesized GnRH neuropeptide. ChINs of the striatum contribute as interneurons to the regulation of cortico-striato-thalamocortical neural pathways. Functions associated with this circuitry include motor control, learning, language, reward, cognitive functioning, and addiction (Fazl et al., 2018). The exact role of GnRH/GnRHR1 signaling in these functions requires clarification. Cholinergic neurons of the nbM which project to the entire cortical mantle, the olfactory tubercle, and the amygdala have been implicated in the control of attention, in the maintenance of arousal, and in learning and memory formation (Koulousakis et al., 2019).

### GnRHR1 signaling may become a therapeutic target to treat cholinergic dysfunctions

Dysfunctions unrelated to the reproductive systems have not been characterized in GnRH deficient patients (Chan, 2011). Future studies will need to clarify alterations of extrahypothalamic GnRH/GnRHR1 signaling in neurodegenerative disorders affecting cholinergic systems. Leading symptoms and cognitive decline in Azheimer’s disease are due to the loss of basal forebrain cholinergic neurons many of which exhibited GnRH immunoreactivity in nbM. Parkinson’s disease (PD) is characterized by motor symptoms such as abnormal involuntary movements, bradykinesia, rigidity, gait, and tremor. Non-motor symptoms often include cognitive impairment, mood disorders, sleep alterations, dysautonomia, anosmia and hallucinations (Perez-Lloret et al., 2016; Tubert et al., 2020). Many of these malfunctions in PD can be explained with the loss of the nigrostriatal dopaminergic input and ameliorated with levodopa. However, gait disorders, cognitive impairment/dementia are most often unresponsive to dopamine precursor treatment. These data indicate involvement of other neurotransmitter systems. In particular, loss of striatal dopamine input causes a local hypercholinergic state in the striatum with consequences reviewed recently (Tubert et al., 2020). This hypercholinergic state explains the success of early PD therapies with *atropa belladonna* derivatives (Goetz, 2011). Although the low efficacy of anticholinergic drugs compared to levodopa and unwanted side effects limit the use of general anticholinergic strategies (Katzenschlager et al., 2003), selective inhibition of striatal ChINs has been proposed recently as a more promising strategy to improve the transmitter balance in dopamine-deprived basal ganglia (Mallet et al., 2019; Tubert et al., 2020). An important physiological mechanism to inhibit acetylcholine release from ChINs is via M2-type (M2 and M4) muscarinic autoreceptors coupled to Gi proteins. Accordingly, deletion of M2-type autoreceptors results in increased striatal acetylcholine release (Bonsi et al., 2008). Autoinhibitory mechanism by muscarinic autoreceptors was found to be lost in PD animal models (Ding et al., 2006). Indeed, our RNA-Seq analysis established that human ChINs contain very high levels of *CHRM2* autoreceptors (**Fig. 4** and **Supplementary File 6**). The receptor transcriptome profile of these cells (**Fig. 4** and **Supplementary File 6**) offers a few alternative mechanisms to inhibit the striatal hypercholinergic state in PD. In particular, selective GnRHR1 agonist treatment which inhibits ChINs in our neonatal mouse model, or induction of *GNRH1* expression in human ChINs may prove to be useful strategies to counteract the hyperactivity of ChINs in PD.

## Conclusions

This study reports discovery and characterization of 150,000-200,000 GnRH-IR neurons which are located in the basal ganglia and the basal forebrain of the adult human brain. These extrahypothalamic GnRH cells represent subsets of previously known cholinergic neurons and synthesize *bona fide* GnRH decapeptide. Unexpectedly, a large subpopulation of hypothalamic GnRH neurons share this cholinergic (ChAT) neurochemistry. Using a neonatal mouse model, GnRHR1 autoreceptor activation reduces the resting membrane potential and electric activity within ChINs of the CPU. Human relevance of these functional data has been shown by the results of RNA-Seq experiments on ChINs and SPNs of the human Pu, which showed that ChINs express *GNRH1*, GnRH biosynthetic enzymes, *GNRHR1* autoreceptors and several inhibitory G-protein isoforms. The role of GnRH/GnRHR1 signaling within extrahypothalamic neuronal circuitries and higher-order functions regulated by GnRH will require clarification.

## Materials and Methods

### Human subjects

Adult human brain tissues were collected from autopsies (N=28) at the 1^st^ Department of Pathology and Experimental Cancer Research, Semmelweis University, Budapest, Hungary. Ethic permissions were obtained from the Regional and Institutional Committee of Science and Research Ethics of Semmelweis University (SE-TUKEB 251/2016), in accordance with the Hungarian Law (1997 CLIV and 18/1998/XII.27. EÜM Decree/) and the World Medical Association Declaration of Helsinki. The demographic data of donors and use of their tissue samples in the different experiments are summarized in **Supplementary File 4**, whereas the most important details of IHC studies are shown schematically in **Supplementary File 5**. The dissected adult tissue blocks were rinsed briefly with running tap water. Then, depending on use, they were either immersion-fixed with buffered paraformaldehyde (PFA) as detailed below or snap-frozen on powdered dry-ice.

### Human fetuses

Fetal tissues (**Supplementary File 4**; #29, 30) were made available in accordance with French bylaws (Good Practice Concerning the Conservation, Transformation, and Transportation of Human Tissue to Be Used Therapeutically, published on December 29, 1998). The studies on human fetal tissue were approved by the French agency for biomedical research (Agence de la Biomédecine, Saint-Denis la Plaine, France, protocol n°: PFS16–002). Non-pathological human fetuses (N=2) were obtained at gestational week 11 (GW11) from pregnancies terminated voluntarily after written informed consent of the parents (Gynaecology Department, Jeanne de Flandre Hospital, Lille, France).

### Mapping and quantitative analysis of extrahypothalamic GnRH neurons in adult brains

Brains (N=3) were cut into ∼15 mm thick coronal slices. The tissue slabs were immersion-fixed in several changes of buffered (0.1 M PBS; pH 7.4) 4% PFA for 21 days and then, infiltrated with 20 % sucrose for 7 days (4 °C). The right hemispheres were isolated and processed to determine the distribution and number of extrahypothalamic GnRH neurons in the nucleus caudatus (Cd), putamen (Pu), globus pallidus (GP), nucleus accumbens (nAcc), bed nucleus of the stria terminalis (BnST) and nucleus basalis of Meynert (nbM). Brain slices were embedded in Jung tissue freezing medium (Leica Biosystems, Nussloch, Germany), snap-frozen on powdered dry ice. Then, 100-μm-thick coronal sections were collected with a Leica SM 2000R freezing microtome into tissue culture plates filled with anti-freeze solution (30% ethylene glycol; 25% glycerol; 0.05 M phosphate buffer; pH 7.4) and stored at −20 °C. Every 24^th^ section between Bregma levels −22.5 and 33.1 (Mai et al., 1997) was immunostained using a well-characterized guinea pig antiserum (#1018) against GnRH decapeptide (Hrabovszky et al., 2011) (**Fig. 1A**). The sections were rinsed in PBS, pretreated with a mixture of 1% H_2_O_2_ and 0.5% Triton X-100 for 30 min and subjected to antigen retrieval in 0.1 M citrate buffer (pH 6.0) at 80 °C for 30 min. To maximize signal, mmunohistochemical incubations were extended: guinea pig anti-GnRH antibodies (#1018; 1:30,000) (Hrabovszky et al., 2011), 5 days; biotinylated donkey anti-guinea pig IgG antibodies (Jackson ImmunoResearch Europe, Cambridgeshire, UK; 1:500), 12 h; ABC Elite reagent (Vector, Burlingame, CA; 1:1000), 4 h. The signal was visualized with nickel-diaminobenzidine (Ni-DAB) chromogen (10 mg diaminobenzidine, 30 mg nickel-ammonium-sulfate and 0.003% H_2_O_2_ in 24 ml Tris-HCl buffer solution (0.05M; pH 8.0). Immunostained sections were mounted on 75 mm X 50 mm microscope sides from 0.3% polyvinyl alcohol, air-dried, dehydrated with 70%, 95% and 100% ethanol (5 min each), cleared with xylenes (2X5 min) and coverslipped with DPX mounting medium (Merck, Darmstadt, Germany).

Anatomical sites to be analyzed separately were identified at each rostro-caudal level (Mai et al., 1997) by macroscopic and microscopic analyses and their borders were marked on the coverslips. Labeled cell bodies were counted in each region with light microscopy and cell numbers were corrected against overcounting (**Supplementary File 1**) using Abercrombie’s correction factor T/(T+h), where T is actual section thickness and h is the average diameter of GnRH neurons along the Z axis (Guillery, 2002). Two Pu sections were used to determine T and h. These sections were processed for the immunofluorescent detection of GnRH neurons with guinea pig anti-GnRH antibodies (#1018; 1:30,000; 12 h), followed by peroxidase-conjugated anti-guinea pig IgG (Jackson ImmunoResearch; 1:250; 2h) and FITC-tyramide (1:1000; 30 min). The sections were embedded into 4% agarose, resectionned with a Leica vibratome perpendicularly to the original section plane. T and h were measured with confocal microscopy to calculate a final correction factor of 0.712. The number of GnRH cells (n) counted in every 24^th^ section of a single hemisphere was first doubled (with the assumption that the distribution of extrahypothalamic GnRH neurons is symmetrical) and then, multiplied by 24 and Abercrombie’s correction factor to estimate the total number of extrahypothalamic GnRH neurons (Σ = n X 2 X 24 X 0.712) in the basal ganglia and the basal forebrain of each brain.

### Immuno-peroxidase detection of extrahypothalamic GnRH neurons using different primary antibodies

Dissected tissue samples (N=10) containing the extrahypothalamic regions of interest were fixed by immersion in freshly-prepared 4% PFA in PBS for 14-21 days at 4 °C. The fixed blocks were trimmed, infiltrated with 20% sucrose for 5 days at 4 °C, placed in a freezing mold, surrounded with Jung tissue freezing medium, snap-frozen on powdered dry ice, and sectioned coronally at 20-30 μm with a freezing microtome (Leica Biosystems). The sections were stored permanently in anti-freeze solution (30% ethylene glycol; 25% glycerol; 0.05 M phosphate buffer; pH 7.4) at −20 °C. Following the pretreatments detailed above, a series of different GnRH and GAP1 antibodies (**Supplementary File 5**) were tested for reactivity with extrahypothalamic GnRH neurons. These included guinea pig (#1018; 1:30,000) (Hrabovszky et al., 2011), rat (#1044; 1:20,000) (Skrapits et al., 2015) and sheep (#2000; 1:1,000) (Skrapits et al., 2015) polyclonal antisera generated in our laboratory against the GnRH decapeptide and the LR1 rabbit GnRH antiserum (1:10,000; gift from Dr. R.A. Benoit) which was reported not to produce specific labeling of extrahypothalamic GnRH neurons in rhesus monkeys (Quanbeck et al., 1997; Terasawa et al., 2001). In addition, a rabbit polyclonal antiserum (MC-2; 1: 5,000) (Culler et al., 1986) to aa 25-53 of hGAP1 (accession: P01148) was used. The signals were detected using biotinylated secondary antibodies (Jackson ImmunoResearch; 1:500; 1h), ABC Elite reagent (Vector; 1:1,000; 1h), and Ni-DAB chromogen and coverslipped with DPX.

### Dual-label immunofluorescence experiments used as a positive control for GnRH labeling

Positive control experiments with immunofluorescence (IF) double-labeling used two sequential rounds of tyramide signal amplification (TSA) to maximize both GnRH signals. The sections were pretreated as above, followed by an additional Sudan Black step (Mihaly et al., 2000) to reduce tissue autofluorescence. Then, a mixture of guinea pig GnRH (#1018; 1:30,000) and rat GnRH (#1044; 1:20,000) primary antibodies were applied to the sections for 16h at 4°C, followed by peroxidase-conjugated anti-guinea pig IgG (Jackson ImmunoResearch; 1:250; 1h) and Cy3-tyramide (Hopman et al., 1998) (diluted 1:1,000 with 0.05 M Tris-HCl buffer/0.005% H_2_O_2_; pH 7.6). Peroxidase was inactivated with 0.5% H_2_O_2_ and 0.1 M sodium azide in PBS for 30 min. Then, the rat GnRH antibodies were reacted with biotin-conjugated secondary antibodies (Jackson ImmunoResearch; 1:500; 1h), ABC Elite reagent (1:1,000, 1h) and FITC-tyramide (Hopman et al., 1998) (diluted 1:1,000 with 0.05 M Tris-HCl buffer/0.005% H_2_O_2_; pH 7.6). The dual-labeled sections were mounted and coverslipped with the aqueous mounting medium Mowiol.

### *In situ* hybridization detection of *GNRH1* mRNA in GnRH neurons of the human putamen

The digoxigenin-labeled antisense probe targeting bases 32-500 of human *GNRH1* mRNA (NM_001083111.2) was transcribed in the presence of digoxigenin-11-UTP (Merck Millipore) in a reaction mixture containing linearized cDNA template (1 µg), 5X transcription buffer (2 µl), 100 mM DTT (1 µl), 10 mM ATP, CTP, and GTP (0.5 µl each), 10 mM digoxigenin-11-UTP (0.5 µl), 1 mM UTP (1 µl), 40 U/µl RNase inhibitor (RNasin; Promega, Madison, WI; 0.5 µl) 20 U SP6 RNA polymerase (Promega; 1 µl). Following a 1-h incubation of the cocktail at 37 °C, a second 20 U aliquot of SP6 RNA polymerase was added and the reaction was allowed to proceed for an additional 1 h. The volume was brought up to 90 µl with nuclease-free water, and the cDNA template was digested for 30 min at 37 °C after the addition of 1 µl DNase I (10 U/µl; Roche Diagnostics, Rotkreuz, Switzerland), 5 µl 1M Tris/HCl buffer (pH 8.0), 1 µl transfer RNA (tRNA; 25 mg/ml), 1 µl 1 M MgCl_2_ and 0.5 µl RNasin (40 U/µl) to the reaction mixture. The cRNA probe was purified using sodium chloride/ethanol precipitation, dissolved in 100 µl of 0.1% sodium dodecyl sulfate, stored at −20 °C and added to the hybridization buffer (50% formamide, 2X SSC, 20% dextran sulfate, 1X Denhardt’s solution, 500 µg/ml yeast tRNA, 50 mM DTT) at a 1:100 dilution (1XSSC = 0.15 M NaCl/0.015 M sodium citrate, pH 7.0). Four-mm-thick putamen blocks were dissected out from the brains (N=5), immersion-fixed in 4% PFA for 48 h and infiltrated with 20% sucrose for 48 h. 20-µm-thick floated sections were prepared with a freezing microtome and processed for combined *in situ* hybridization (ISH) detection of *GNRH1* mRNA and IF detection of GnRH peptide. First, the sections were acetylated with 0.25% acetic anhydride in 0.9% NaCl/0.1 M triethanolamine-HCl for 10 min, rinsed in 2X SSC for 2 min, treated sequentially with 50%, 70%, and 50% acetone (5 min each), rinsed with 2X SSC, and hybridized overnight in microcentrifuge tubes containing the hybridization solution. Non-specifically bound probes were digested with 20 µg/ml ribonuclease A (Merck; dissolved in 0.5 M NaCl/10 mM Tris-HCl/1 mM EDTA; pH 7.8) for 60 min at 37 °C, followed by a 60-min-stringent treatment (55 °C in 0.1XSSC solution for) to reduce background. The floated sections were rinsed briefly with 100 mM maleate buffer (pH 7.5) and blocked for 30 minutes against non-specific antibody binding with 2% blocking reagent (Merck) in maleate buffer. To detect the hybridization signal, the sections were incubated overnight at 4 °C in digoxigenin antibodies conjugated to horseradish peroxidase (anti-digoxigenin-POD; Fab fragment; 1:100; Roche), rinsed in TBS (0.1 M Tris-HCl with 0.9% NaCl; pH 7.8) and then, reacted with Cy3-tyramide (Hopman et al., 1998) (diluted 1:1,000 with 0.05 M Tris-HCl buffer/0.005% H_2_O_2_; pH 7.6) for 30 min. Peroxidase was inactivated with 0.5% H_2_O_2_ and 0.1 M sodium azide in PBS for 30 min. Subsequently, GnRH immunoreactivity was detected with guinea pig anti-GnRH (#1018; 1:30,000) primary antibodies (16 h at 4 °C), biotin-conjugated secondary antibodies (Jackson ImmunoResearch; 1:500; 1h), ABC Elite reagent (1:1,000, 1h) and FITC-tyramide (Hopman et al., 1998) (diluted 1:1,000 with 0.05 M Tris-HCl buffer/0.005% H_2_O_2_; pH 7.6).

### DiI-labeling of putamen sections to study GnRH cell morphology

Combined immunofluorescent detection of peptidergic neurons and their Golgi-like cell membrane labeling with the lipophilic dye DiI using a Gene Gun was adapted to studies of human extrahypothalamic GnRH neurons from our recently reported procedure (Takacs et al., 2018). A 4-mm-thick tissue block was dissected from the Pu and immersion-fixed lightly with freshly prepared 2% PFA in 0.1 M PBS (pH 7.4) for 14 days (4 °C). 100-µm-thick coronal sections were prepared with a Leica VTS-1000 Vibratome (Leica Biosystems) and stored in PBS/0.1% sodium azide at 4 °C before use. The sections were pretreated with a mixture of 1% H_2_O_2_ and 0.5% Tween 20 for 30 min, followed by epitope retrieval with 0.1 M citrate buffer (pH 6.0) at 80 °C for 30 min. Then, sequential incubations were carried out in the guinea pig GnRH antibodies (#1018; 1:30,000) for 4 days, peroxidase-conjugated anti-guinea pig antibodies (Jackson ImmunoResearch Laboratories; 1:250) for 4 h, and finally, FITC-tyramide (diluted 1:1,000 with 0.05 M Tris-HCl buffer/0.005% H_2_O_2_; pH 7.6; 30 min) prepared (Hopman et al., 1998) and used (Takacs et al., 2018) as reported. Methods to prepare and deliver DiI-coated tungsten particles with a Helios Gene Gun (Bio-Rad, Hercules, CA) were adapted from published procedures (Seabold et al., 2010; Staffend et al., 2011). Sections of the Pu were transferred into 12-well tissue culture plates containing PBS. The buffer was removed with a pipette and diolistic labeling was carried out using a 40-mm spacer and a 120-150 pounds per square inch (PSI) helium pressure, which resulted in random-labeling of cells, including 12 GnRH-IR neurons. Labeled sections were rinsed in PBS/0.1% sodium azide/0.2% EDTA and the lipophilic dye was allowed to diffuse along the cytoplasmic membranes for 24 h at 4 °C. The sections were coverslipped with Mowiol to study the Golgi-like DiI labeling of the randomly hit GnRH neurons.

### Dual-label immunofluorescence experiments to colocalize choline acetyltransferase with GnRH

Sections from striatal (N=4) and hypothalamic (N=7) samples were rinsed in PBS followed by a mixture of 1% H_2_O_2_ and 0.5% Triton X-100 for 30 min, and then, subjected to antigen retrieval in 0.1M citrate buffer (pH=6.0) at 80 °C for 30 min and Sudan Black pretreatment. GnRH neurons were detected using sequentially guinea pig GnRH antibodies (#1018; 1:30,000; 16h; 4 °C), peroxidase-conjugated anti-guinea pig IgG (Jackson ImmunoResearch; 1:250; 1h) and FITC-tyramide (Hopman et al., 1998) (diluted 1:1,000 with 0.05 M Tris-HCl buffer/0.005% H_2_O_2_; pH 7.6). Peroxidase was inactivated with 0.5% H_2_O_2_ and 0.1 M sodium azide in PBS for 30 min. Then, ChAT neurons were detected using goat anti-ChAT antibodies (AB144P; Merck; 1:2,000) (Yonehara et al., 2011), biotinylated secondary antibodies (donkey anti-goat IgG; Jackson ImmunoResearch; 1:500), ABC Elite reagent (Vector) and Cy3-tyramide (diluted 1:1,000 with 0.05M Tris-HCl buffer, pH 7.6, containing 0.005% H_2_O_2;_ 30 min) (Hopman et al., 1998). The dual-labeled sections were mounted on slides, coverslipped with Mowiol and analyzed with confocal microscopy. Confocal Z-stacks were prepared from each region and analyzed to determine the percentage of GnRH neurons showing ChAT immunoreactivity and, *vice versa*.

### Dual-imunofluorescence studies of fetal tissues

The fetuses (N=2) were fixed by immersion in 4% buffered PFA at 4 °C for 5 days. The tissues were then cryoprotected in PBS containing 30% sucrose at 4°C overnight, embedded in Tissue-Tek OCT compound (Sakura, Finetek), frozen in dry ice and stored at −80 °C until sectioning. Frozen samples were cut serially at 20 μm with a cryostat (Leica Biosystems) and immunolabeled, as described previously (Casoni et al., 2016), with polyclonal goat anti-ChAT (AB144P; Merck; 1:150) and guinea pig anti-GnRH antibodies (#1018; 1:10,000), in a solution containing 10% normal donkey serum and 0.3% Triton X100 at 4 °C for 3 days. 3 x 10 min washes in 0.01 M PBS were followed by incubations in AF568-conjugated donkey anti-goat (Invitrogen; 1:400) and AF488-conjugated donkey anti-guinea pig (Jackson ImmunoResearch; 1:400) antibodies for 1h each. The section were counterstained with Hoechst (1:1,000) and coverslipped with Mowiol.

### RNA-sequencing

#### Reagents for RNA-Seq

For all experiments, nuclease-free water was used and reagents were of molecular biology grade. Work surfaces and equipments were cleaned with RNaseZAP.

#### Section preparation for RNA-seq experiments

After dissection, tissue samples from the Pu of two subjects (#21 and 22) were snap-frozen in −40 °C isopentane precooled with a mixture of dry ice and ethanol. Then, 20 µm-thick coronal sections were cut with a Leica CM1860 UV cryostat (Leica Biosystems. Wetzlar, Germany), collected onto PEN membrane glass slides (Membrane Slide 1.0 PEN, Carl Zeiss, Göttingen, Germany), air-dried for 5 min in the cryostat chamber, and fixed with a mixture of 2% PFA, 0.1% diethyl pyrocarbonate, 1% sodium acetate and 70% ethanol (10 min). After brief rehydration (RNase-free water 2 min), sections were stained with 0.5% cresyl violet solution (1 min), rinsed in RNase-free water and dehydrated again in 70, 96 and 100% ethanol (30 sec each). The slides were kept at −80 °C in clean slide mailers containing silica gel desiccants until further processing.

#### Laser-capture microdissection

Slides were placed into the slide holder of the microscope and 300 cholinergic interneurons (ChINs) were microdissected by LCM using a PALM Microbeam system (Zeiss). The cells were pressure-catapulted from the object plane into 0.5 ml tube caps (Adhesive Cap 200, Zeiss) with a single laser pulse using a 40x objective lens. A second control cell pool was prepared from 600 medium sized neurons most of which corresponded to SPNs. The mean profile areas of ChINs and SPNs were 674.76 µm^2^ and 161.22 µm^2^, respectively. The LCM caps were stored at −80 °C until RNA extraction.

#### RNA extraction, RNA-seq library preparation and sequencing

The Arcturus Paradise Plus RNA Extraction and Isolation Kit (Thermofisher, Waltham, MA, USA) was used to isolate total RNA according to the manufacturers protocol. Samples collected from control sections of the two brains showed RNA integrity numbers (RINs) of 5.7 and 4.1, respectively, as determined using Bioanalyzer Eukaryotic Total RNA Pico Chips (Agilent, Santa Clara, CA, USA). RNA samples were converted to RNA-seq libraries with the TruSeq Stranded Total RNA Library Preparation Gold kit (Illumina, San Diego, CA, USA). This kit was reported to reliably and reproducibly generate libraries from 1-2 ng input RNA (Schuierer et al., 2017). The manufacturer’s protocol was followed, except for the use of 17, instead of 16, cycles of amplification for adaptor-ligated DNA fragment enrichment. Single-end sequencing was performed on Illumina NextSeq500 instrument using the Illumina NextSeq500/550 High Output kit v2.5 (75 Cycles).

#### Bioinformatics

After quality check with FastQC, raw reads were cleaned by trimming low-quality bases by Trimmomatic 0.39 (settings: LEADING:3, TRAILING:3, SLIDINGWINDOW:4:30, MINLEN:50). The prepared reads were mapped to the GRCh38.p13 human reference genome using STAR (v 2.7.3a) (Dobin et al., 2013) with an average overall alignment rate of 68.4% (s.d. = 9.9%). Gene level quantification of read counts based on human genome with Ensembl (release 99) (Yates et al., 2020) annotation was performed by featureCounts (subread v 2.0.0) (Liao et al., 2014), with a mean of 30.4% (s.d. = 9.7%) of mapped reads assigned to genes in the case of the four samples. The raw read counts per genes were normalized and processed further in R (R2020) with the package DESeq2 (Love et al., 2014) and edgeR (McCarthy et al., 2012). For feature annotation, the R package KEGGREST (Dan Tenenbaum, KEGGREST: Client-side REST access to KEGG. R package version 1.26.1.; <u>2019</u>) and the PANTHER database (v. 15.0) (Thomas et al., 2003) were used.

### High Performance Liquid Chromatography-tandem mass spectrometry (HPLC-/MS/MS)

Brain tissue specimens were snap-frozen and kept at −80 °C. ∼10-60 mg samples (**Supplementary File 4**) were microdissected in a −20 °C cryostat chamber from the MBH, Pu, Cd and Cl. After addition of the extraction solution containing 1% acetic acid and Complete Mini protease inhibitor cocktail (Roche, Basel, Switzerland) in 1:2 w/v proportion, samples were homogenized using an ultrasonic sonotrode. The homogenates were mixed with double volume acetonitrile and centrifuged to produce protein-free supernatants. Separation of 10 µl samples was carried out by HPLC (Perkin Elmer Series 200) using gradient elution on a Luna Omega Polar C18 50×3 mm, 3 μm column (Phenomenex, Torrance, CA, USA). Acetonitrile and 0.1% formic acid were applied for gradient elution with the flow rate of 500 µl/min. Acetonitrile increased from 10% to 40% in 3 min, and this was maintained for 0.5 min. The initial 10% was reached in 0.5 min and maintained for 2 min. Analytes were detected using a triple quadrupole MDS SCIEX 4000 Q TRAP mass spectrometer (Applied Biosystems) in positive multiple reaction monitoring mode (MRM transitions: GnRH: 592.1 → 249.3, GnRH1-5: 671.2 → 159.1). Peak areas were integrated with Analyst 1.4.2 software (Sciex, Framingham, MA, USA), and concentrations were calculated using matrix-matched calibration.

### Animals

Experiments involving genetically modified male mice were carried out in accordance with the Institutional Ethical Codex, Hungarian Act of Animal Care and Experimentation (1998, XXVIII, section 243/1998) and the European Union guidelines (directive 2010/63/EU), and with the approval of the Institutional Animal Care and Use Committee of the Institute of Experimental Medicine. The animals were housed under standard conditions (lights on between 06.00 and 18.00 h, temperature 22±1 °C, chow and water *ad libitum*) and all measures were taken to minimize potential stress or suffering during sacrifice and to reduce the number of animals to be used. i) ChAT-Cre/zsGreen mice were generated by crossing ChAT-IRES-Cre knock-in mice (Jackson Laboratory, Bar Harbor, ME; RRID: IMSR_JAX:006410) with the Ai6(RCL-ZsGreen) indicator strain (The Jackson Laboratory, JAX No. 007906). Mice used for the experiments were heterozygous for both the Cre and the indicator gene alleles. ii) GnRH-GFP transgenic mice which selectively express enhanced green fluorescent protein in GnRH neurons were generated in the laboratory of Dr. S.M. Moenter (Suter et al., 2000) and maintained as a homozygous colony for the transgenes. iii) The gad65-GFP (FVB.129 Tg gad65-gfp) transgenic mice were generated by Ferenc Erdélyi and Gábor Szabó and characterized elsewhere (Lopez-Bendito et al., 2004). Their colony was maintained as heterozygous for the transgene and positive offprings were identified by direct fluorescence visualization.

### Perfusion-fixation and section preparation for anatomical studies

Male GnRH-GFP transgenic mice (N=9) were anesthetized between 0900 and 1100 h with a cocktail of ketamine (25 mg/kg), xylavet (5 mg/kg) and pipolphen (2.5 mg/kg) in saline, and then, perfused transcardially with 4% PFA in 0.1 M PBS (pH 7.4). The brains were removed, infiltrated with 20% sucrose overnight and snap-frozen on powdered dry ice. 25-μm-thick coronal sections were collected from the CPU with a freezing microtome and stored at −20 °C in antifreeze solution. This region in adults corresponded to Atlas plates 20-30 of Paxinos (Bregma levels 1.34-0.14 mm) (Paxinos et al., 2001).

### Simultaneous visualization of GnRH-GFP fluorescence and ChAT-immunoreactivity

Floating sections of the CPU were pretreated with 0.5% H_2_O_2_ and 0.2% Triton X-100. Cholinergic neurons were detected with the AB144P goat ChAT antiserum (1:2,000) and TSA, as described for human studies. The sections were mounted on slides, coverslipped with Mowiol and analyzed with confocal microscopy. Expression of the GnRH-GFP trangene was shown by the green fluorescence of scattered CPU neurons.

### Light microscopy

Representative light microscopic images were prepared with an AxioCam MRc 5 digital camera mounted on a Zeiss AxioImager M1 microscope, using the AxioVision 4.6 software (Carl Zeiss, Göttingen, Germany).

### Confocal microscopy

Fluorescent signals were studied with a Zeiss LSM780 confocal microscope. High-resolution images were captured using a 20×/0.8 NA objective, a 0.6–1× optical zoom and the Zen software (CarlZeiss). Different fluorochromes were detected with laser lines 488 nm for FITC and AF488 and 561 nm for Cy3. Emission filters were 493–556 nm for FITC and AF488 and 570–624 nm for Cy3. To prevent emission crosstalk between the fluorophores, the red channel was recorded separately from the green one (“smart setup” function). To illustrate the results, confocal Z-stacks (Z-steps: 0.85-1 μm, pixel dwell time: 0.79-1.58 μs, resolution: 1024×1024 pixels, pinhole size: set at 1 Airy unit) were merged using maximum intensity Z-projection (ImageJ). The final figures were adjusted in Adobe Photoshop using the magenta-green color combination and saved as TIF files.

Fetal sections were examined using an Axio Imager.Z1 ApoTome microscope (Carl Zeiss, Germany) equipped with a motorized stage and an AxioCam MRm camera (Zeiss). For confocal observation and analyses, an inverted laser scanning Axio observer microscope (LSM 710, Zeiss) with an EC Plan NeoFluorÅ∼100/1.4 numerical aperture oil-immersion objective (Zeiss) was used (Imaging Core Facility of IFR114, of the University of Lille, France).

### Brain slice preparation for electrophysiological recordings

Brain slices of the different transgenic mice were prepared as described earlier (Farkas et al., 2010) and used to record PW1 (postnatal day 4-7) GnRH-GFP (n=13), PW1 ChAT-Cre/zsGreen (n=41), adult ChAT-Cre/zsGreen (n=10), and PW1 GAD65-GFP (n=16) neurons. Briefly, the mice were decapitated in deep inhalation anesthesia with Isoflurane. The brains were immersed in ice-cold low-Na cutting solution bubbled with carbogen (mixture of 95% O_2_ and 5% CO_2_). The cutting solution contained the following (in mM): saccharose 205, KCl 2.5, NaHCO_3_ 26, MgCl_2_ 5, NaH_2_PO_4_ 1.25, CaCl_2_ 1, glucose 10. CPU blocks were dissected, and 200-μm-thick coronal slices were prepared with a VT-1000S vibratome (Leica Biosystems) in ice-cold oxygenated low-Na cutting solution. The slices were transferred into carbogenated artificial cerebrospinal fluid (aCSF) containing in mM: NaCl 130, KCl 3.5, NaHCO_3_ 26, MgSO_4_ 1.2, NaH_2_PO_4_ 1.25, CaCl_2_ 2.5, glucose 10, and allowed to equilibrate for 1 h while temperature was allowed to drop slowly from 33 °C to room temperature.

Recordings were carried out in carbogenated aCSF at 33 °C using Axopatch-200B patch-clamp amplifier, Digidata-1322A data acquisition system, and pCLAMP 10.4 software (Molecular Devices Co., Silicon Valley, CA, USA). The patch electrodes (OD = 1.5 mm, thin wall; WPI, Worcester, MA, USA) were pulled with a Flaming-Brown P-97 puller (Sutter Instrument Co., Novato, CA, USA). Neurons were visualized with a BX51WI IR-DIC microscope (Olympus Co., Tokyo, Japan). GnRH-GFP, ChAT-Cre/zsGreen and GAD65-GFP neurons showing green fluorescence were identified by brief illumination at 470 nm using an epifluorescent filter set.

Whole-cell patch-clamp measurements started with a control recording (2 min). Then, a single bolus of GnRH (final 1.2 µM) was pipetted into the aCSF-filled measurement chamber and the recording continued for 12 min. Pretreatment of slices with the GnRHR1 antagonist Antide (100 nM) or the voltage-gated Na-channel inhibitor tetrodotoxin (TTX, 660 nM) started 10 min before GnRH was added to the aCSF and these inhibitors were present in the aCSF throughout the recording. To block GPCRs in the recorded neurons, Guanosine 5′-[β-thio]diphosphate trilithium salt (GDP-β-S, 2 mM) was added to the intracellular pipette solution. To allow the intracellular milieu to reach equilibrium, the recording was only started 15 min after achieving the whole-cell patch clamp configuration. Each neuron served as its own control when drug effects were evaluated.

### Whole-cell patch clamp experiments

The action potentials (APs) and resting potentials (V_rest_) were recorded in current-clamp mode. V_rest_ was measured at 0 pA. Most of the neurons were silent at 0 pA. Therefore, APs were triggered with a 15 min-long 10 pA depolarizing current pulse throughout the recording. Intracellular pipette solution contained (in mM): HEPES 10, KCl 140, EGTA 5, CaCl_2_ 0.1, Mg-ATP 4, Na-GTP 0.4 (pH 7.3 with NaOH). The resistance of the patch electrodes was 2–3 MΩ. Spike-mediated transmitter release was blocked in some experiments by adding the voltage-sensitive Na-channel inhibitor TTX (660 nM, Tocris) to the aCSF 10 min before V_rest_ was recorded.

### Drugs

Extracellularly used drugs: GnRH decapeptide (1.2 µM, Merck); GnRH-R antagonist Antide (100 nM; Bachem, Bubendorf, Switzerland); voltage-gated Na-channel inhibitor TTX (660 nM, Tocris). Intracellularly applied drug: the membrane impermeable G-protein blocker GDP-β-S (2 mM, Merck) (Farkas et al., 2016; McDermott et al., 2011).

### Statistical analysis

To minimize sampling bias in electrophysiological studies, animals from the same litter were used for different experiments and slices from the same animals were randomized between treatments. Recordings were stored and analyzed off-line. Event detection was performed using the Clampfit module of the PClamp 10.4 software (Molecular Devices Co., Silicon Valley, CA, USA).

Firing rates were calculated from the number of APs in the given recording time (3 min or 12 min). All experiments were self-controlled. Frequencies and V_rests_ following treatments were expressed as percentages of the untreated control periods. Two-tailed Student’s *t*-tests were applied to assess treatment effects which were considered significant at *p*< 0.05. Treatment groups were characterized with the mean ± standard error of mean (SEM) and compared with one-way ANOVA with repeated measurements followed by Tukey’s test.

## Acknowledgements

The research leading to these results has received funding from the National Science Foundation of Hungary (K128317 to E.H.; PD125393 and PD134837 to K.S.), the Hungarian Brain Research Program (2017-1.2.1-NKP-2017-00002 to E.H.), the Institut National de la Santé et de la Recherche Médicale, Inserm, France (grant number U1172) to P.G. and V.P., and from the Agence Nationale de la Recherche, France (grant number ANR-19-CE16-0021-02 to P.G.). We thank the midwives of the Gynecology Department, Jeanne de Flandre Hospital of Lille (Centre d’Orthogénie), France, for their kind assistance and support; M Tardivel and A Bongiovanni (BICeL core facility of Lille, Univ. Lille, CNRS, Inserm, CHU Lille, Institut Pasteur de Lille, US 41-UMS 2014-PLBS, F-59000 Lille, France) for expert technical assistance. The authors acknowledge support of the Inserm Cross-Cutting Scientific Program (HuDeCA). The research carried out at BME has been supported by the NRDI Fund (TKP2020 IES, Grant No. BME-IE-BIO) based on the charter of bolster issued by the NRDI Office under the auspices of the Ministry for Innovation and Technology. The authors are grateful to Dr. R.A Benoit for the LR-1 antiserum and to Dr. Suzanne M. Moenter for the GnRH-GFP mice.

## Author Contributions

**Conceptualization**, K.S., M.S., B.G., C.V., N.S., S.P., V.P., P.G., E.H.; **Methodology**, K.S., M.S., I.F., B.G., S.T., É.R., V.V., G.R., A.M., N.S., S.P., B.T., F.E., G.S., M.D.C., P.G., E.H.; **Investigation**, K.S., M.S., I.F., B.G., S.T., É.R., V.V., N.S., B.T., C.A., L.C., P.G., E.H.; **Writing—editing,** M.S., I.F., N.S., B.T., V.P., P.G., E.H.; **Funding acquisition and supervision**, K.S., V.P., P.G., E.H.

## Competing Interests

The authors declare no competing interests.

## Data availability

The data that support the findings of this study are available from the corresponding author upon reasonable request. RNA sequencing files will be available in BioProject with the accession number PRJNA680536 (release date: 2021-12-24).

## Code availability

Scripts will be available upon request at https://github.com/solymosin/PRJNA680536_ms_codes

## SOURCE DATA FILES

Figure 3 – Source Data

## SUPPLEMENTARY FILES

**Supplementary File 1. Cell numbers determined with light microscopic analysis requires compensation for overcounting using Abercrombie’s correction factor.** Total extrahypothalamic GnRH cell numbers were determined in three brains (#1-3) by counting GnRH-immunoreactive neurons in every 24^th^ 100-µm-thick coronal section of a single hemisphere. This count was multiplied by 24 and then, doubled, with the assumption that neurons are distributed evenly between right and left hemispheres. **A**: When section thickness (T) and cell diameter (h) along the Z axis are close to each other, a relatively high proportion of cells visualized in section 2 will be transsected (asterisks). Note that taking into account the transsected neurons at both surfaces of section 2 causes considerable over-counting within the tissue volume. This systematic error can be corrected by using Abercrombie’s correction factor calculated from the actual section thickness (T) and the mean diameter of uncut GnRH neurons (h) along the Z axis (schemas based on Guillery et al., 2002) (Guillery, 2002). **B**: To determine T and h, GnRH neurons were detected in floating section of the putamen. Then, the sections were embedded into 2% agarose and recut perpendicular to the original section plane with a Vibratome, mounted on glass slides, coverslipped and analyzed with confocal microscopy. Abercrombie’s correction factor obtained with this approach for the putamen was 0.712. Scale bar: 100 µm.

**Supplementary File. 2. Combined evidence from immunohistochemistry, *in situ* hybridization and high prerformance liquid chromatography-tandem mass spectrometry indicates that extrahypothalamic GnRH neurons synthesize *bona fide* GnRH decapeptide derived from the *GNRH1* transcript. A**: A series of different primary antisera against the human GnRH-associated peptide (hGAP1) or GnRH recognize immunoreactive neurons in the human putamen (Pu) using immunohistochemistry (IHC). Such antibodies include the LR1 rabbit primary antiserum which was reported previously not to label extrahypothalamic GnRH neurons in monkeys. **B**: Positive control with the combined use of two GnRH antibodies from different host species for dual-immunofluorescence (IF) experiments provides evidence that the antibodies detect the same neuronal elements. **C**: Non-isotopic *in situ* hybridization (ISH)/IF dual-labeling studies reveal that GnRH-immunoreactive neurons express *GNRH1* mRNA, indicating that extrahypothalamic GnRH is a *GNRH1* gene product. **D**: As illustrated in representative chromatograms, high performance liquid chromatography followed by tandem mass spectrometry (HPLC-MS/MS) detects *bona fide* GnRH decapeptide in tissue extracts from the mediobasal hypothalamus (MBH), putamen (Pu) and nucleus caudatus (Cd), but not the claustrum (Cl). **E**: The GnRH1-5 degradation product is present in the Pu and Cd and undetectable in the MBH and Cl. **F**: Quantitative analysis reveals the highest tissue concentrations of GnRH in the MBH, somewhat lower levels in the Pu and the Cd, and no detectable GnRH decapeptide signal in the Cl. Note that tissue concentrations of GnRH1-5 in the Pu and the Cd are 3-4 times higher than those of GnRH1-5. Scale bar: 50 µm.

**Supplementary File 3. The GnRH-GFP transgene is expressed transiently in the caudate-putamen of neonatal mice. A**: Postnatal week 1 (PNW1) mice exhibit transient green fluorescent protein (GFP) fluorescence in the caudate-putamen (CPU; green) of GnRH-GFP transgenic mice (Suter et al., 2000). The cholinergic marker choline acetyltransferase (ChAT; magenta) is not detectable yet with immunohistochemistry at this age. High-power image shows a GnRH-GFP neuron from the framed region. **B**: PNW2 mice exhibit the GFP signal and also express immunoreactivity to ChAT in the CPU. GnRH-GFP fluorescence occurs selectively within ChAT-immunoreactive cholinergic neurons (high-power inset). **C**: The ChAT signal becomes much stronger by PNW4. By this time the GnRH-GFP fluorescent signal fades away from cholinergic cells (high-power inset). Scale bar: 50 µm, and 25 µm in insets.

**Supplementary File 4. Demographic information about the donors and use of tissue specimens in different experiments.** ChAT, choline acetyltransferase, ChINs, cholinergic interneurons; GW11. gestational week 11; IF, immunofluorescence; IHC, immunohistochemistry; ISH, *in situ* hybridization; HPLC-MS/MS, high performance liquid chromatography-tadem mass spectrometry; PMI, *postmortem* interval; RIN, RNA integrity number; RNA-Seq, RNA sequencing; SPNs, medium spiny projection neurons.

**Supplementary File 5. Basic data on antibody specification, concentration and previous characterization immunohistochemical reagents and applied detection methods.** ChAT, choline acetyltransferase; GAP1, GnRH-associated peptide-1; GnRH, gonadotropin-releasing hormone; IF, immunofluorescence; IHC, immunohistochemistry; TSA, tyramide signal amplification.

**Supplementary File 6. Detailed receptor expression profile of ChINs and SPNs from the human putamen.** (Numeric values in cpm.)

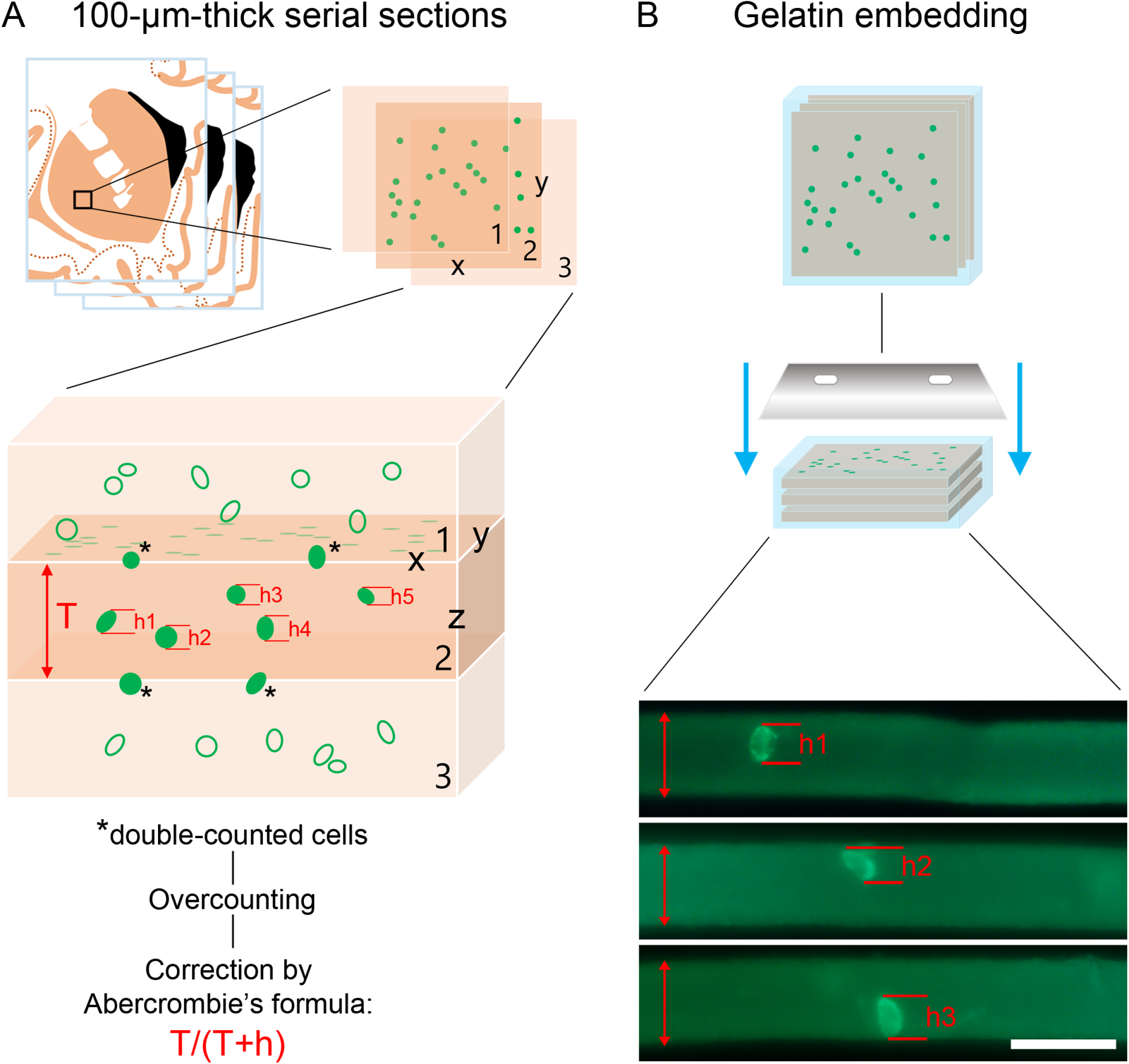

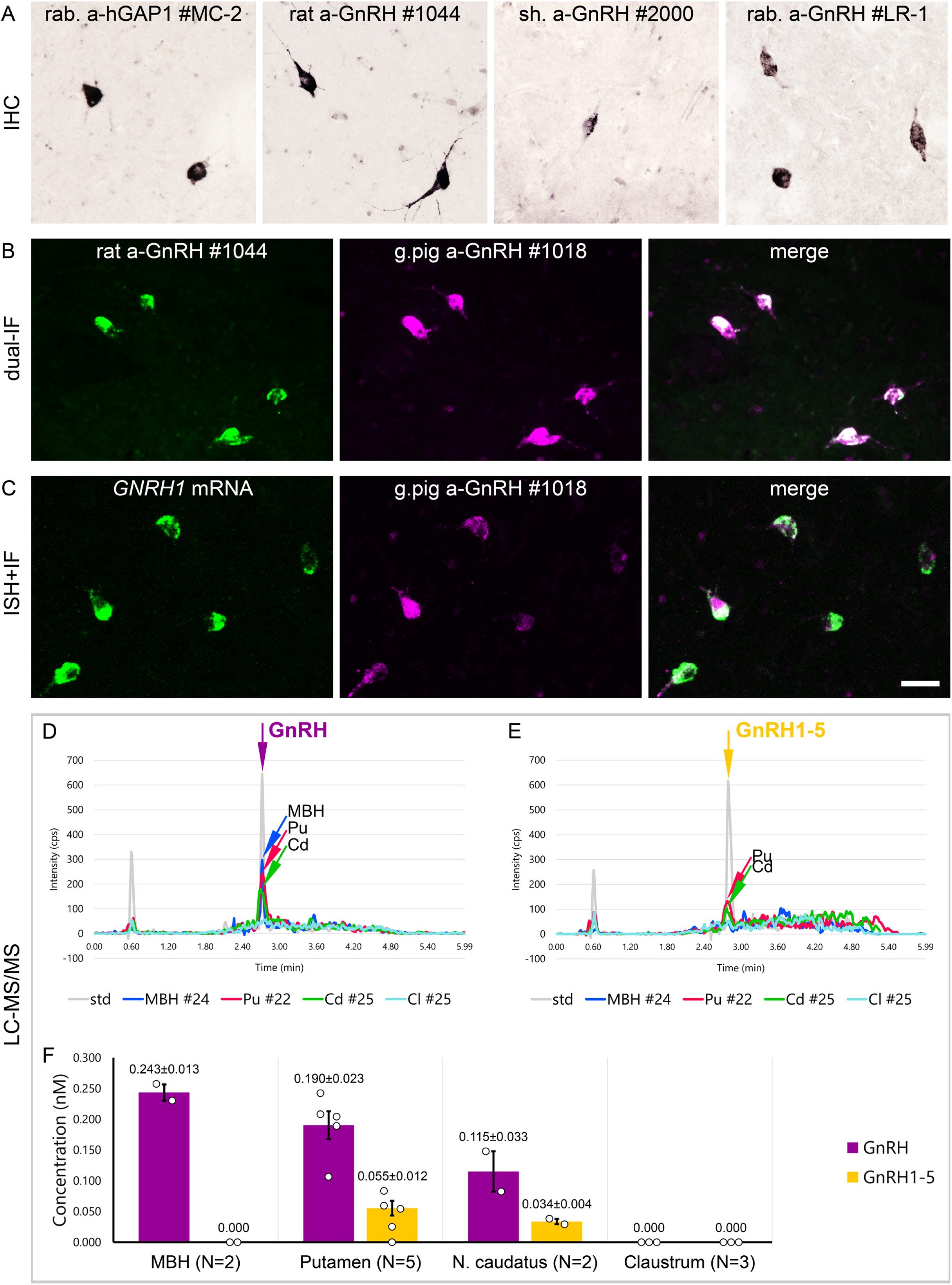

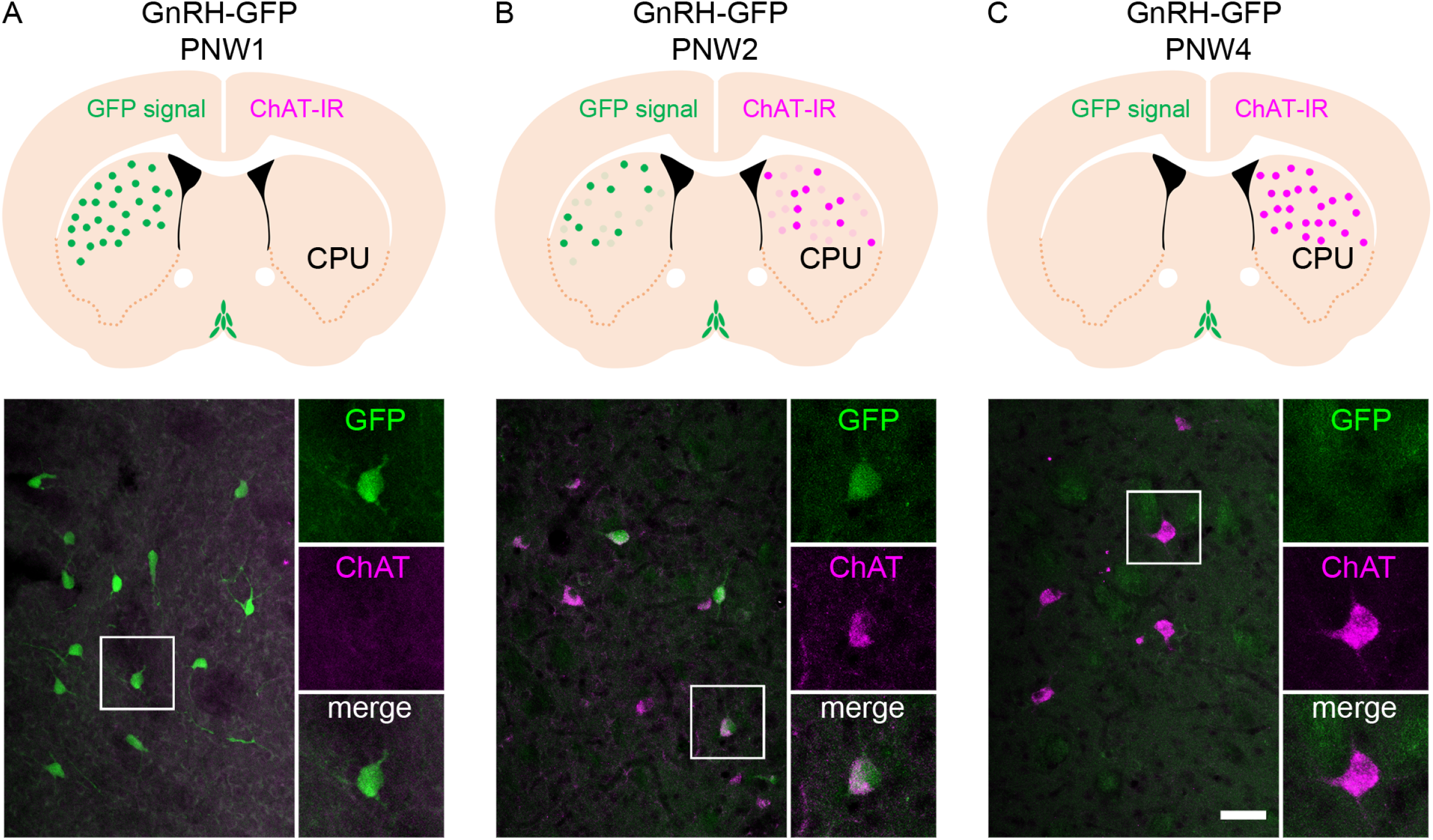

